# Covariant Fitness Clusters Reveal Structural Evolution of SARS-CoV-2 Polymerase Across the Human Population

**DOI:** 10.1101/2022.01.07.475295

**Authors:** Chao Wang, Nadia Elghobashi-Meinhardt, William E. Balch

**Affiliations:** Department of Molecular Medicine, Scripps Research, La Jolla, California, 92037, USA; Department of Physical and Theoretical Chemistry, Technische Universität Berlin, 10623 Berlin, Germany.

## Abstract

Understanding the fitness landscape of viral mutations is crucial for uncovering the evolutionary mechanisms contributing to pandemic behavior. Here, we apply a Gaussian process regression (GPR) based machine learning approach that generates spatial covariance (SCV) relationships to construct stability fitness landscapes for the RNA-dependent RNA polymerase (RdRp) of SARS- CoV-2. GPR generated fitness scores capture on a residue-by-residue basis a covariant fitness cluster centered at the C487-H642-C645-C646 Zn^2+^ binding motif that iteratively evolves since the early phase pandemic. In the Alpha and Delta variant of concern (VOC), multi-residue SCV interactions in the NiRAN domain form a second fitness cluster contributing to spread. Strikingly, a novel third fitness cluster harboring a Delta VOC basal mutation G671S augments RdRp structural plasticity to potentially promote rapid spread through viral load. GPR principled SCV provides a generalizable tool to mechanistically understand evolution of viral genomes at atomic resolution contributing to fitness at the pathogen-host interface.

## Introduction

Coronavirus disease 2019 (COVID-19) (1, 2), caused by the severe acute respiratory syndrome coronavirus 2 (SARS-CoV-2) (3, 4), has resulted in more than 299 million cases and led to more than 5.4 million deaths globally (5). SARS-CoV-2 is a positive-sense single-stranded RNA virus with a ∼30 kb genome that has an estimated variation rate of 1.12 × 10^-3^ mutations per site-year (6). More than 0.15 million unique genetic mutations have been identified for SARS- CoV-2 based on more than 3.7 million SARS-CoV-2 genome sequences (7–12). Understanding the structural and functional impact of these mutations and linking this information to their spread in the human population are crucial in revealing the mechanism of adaptation leading to fitness of SARS-CoV-2 in the human host environment (13, 14) as well as in guiding the surveillance and management of the COVID-19 pandemic.

Our current understanding of SARS-CoV-2 mutations principally comes from studies on the spike protein given its central role in mediating the entry into the host cells and serving as the primary target for neutralizing antibodies (15–17). For example, the D614G mutation in the spike protein found in dominant lineages has been shown to increase the infection rate and transmission for SARS-CoV-2 (18–22). Moreover, N501Y, E484K and L452R mapping to the receptor binding domain of the spike protein and emerging in <u>v</u>ariants <u>o</u>f <u>c</u>oncern (VOC) (e.g., Alpha (B.1.1.7), Beta (B.1.351), Gamma (P.1), Delta (B.1.617.2/AY.*), and Omicron (B.1.1.529)) have drawn attention due to their potential roles in mediating the host cell receptor binding and/or in enabling immune escape (16, 17). Besides the spike protein, SARS-CoV-2 encodes ∼29 additional proteins including 16 <u>n</u>on-<u>s</u>tructural <u>p</u>roteins (nsp), four structural proteins and other accessory proteins (23, 24). Each protein interacts with a set of host proteins to establish the many aspects of the SARS-CoV-2 life cycle (23, 25). However, beyond the well-studied spike protein, little is known about the impact of mutations on the function and structure of other SARS-CoV-2 proteins responsible for VOC linked surges driving the pandemic (15, 26).

The replication and transcription of SARS-CoV-2 genome are managed by the viral RNA- dependent RNA polymerase (RdRp) protein complex, which is comprised of the catalytic subunit nsp12 and the cofactors nsp7 and nsp8 (27–30). Nsp12 alone has little activity and its binding with nsp7 and two nsp8 subunits (referred to herein as nsp8-1 and nsp8-2) is necessary for the polymerase function (27, 31). Nsp12 is the target of Remdesivir, a nucleoside analog that inhibits RdRp activity through chain termination (28, 32, 33), and more recently, the nucleoside analog molnupiravir that triggers lethal mutagenesis (34–36). Both are approved by U.S. Food and Drug Administration (FDA) to treat COVID-19 patients, although clinical trials using a broad population spanning multiple countries found no significant effect of Remdesivir in preventing disease progression (44). Characterizing the functional and structural fitness of RdRp mutations during SARS-CoV-2 transmission in the context of the world population is important for understanding the physiological mechanisms responsible for its ability to adapt to the human host environment(s). These mechanisms that potentially impact viral replication and viral load are important for the development of novel therapeutic strategies to limit viral transmission and pathology.

In the evolution of RNA viruses, an increase in mutation frequency can arise from genetic drift due to random events, or from positive selection according to Darwinian principles (45). Identifying the corresponding functional and structural selection processes driving fitness in response to variation is challenging, particularly for those directing complex pathogen-host relationships (26, 46, 47). Here, we assess the impact of mutations on the structural thermodynamic stability of RdRp through free energy computation reflecting physical-chemical features contributing to fold function. These values are then linked to virus spread in the human population using a <u>G</u>aussian <u>p</u>rocess <u>r</u>egression (GPR) based machine learning approach termed variation spatial profiling (VSP) (48–50). VSP is based on the principle of spatial covariance (SCV) that connects the sequence and phenotype information of a sparse collection of mutations found in the population to pinpoint and project the critical residue-residue relationships shaped through evolution driving function (48). GPR also generates robust uncertainty for each prediction (51–54), providing a rigorous tool to assess the probability of function of a residue in the context of other evolving residues in response to the local environment (48).

Using GPR, we construct ‘fitness landscapes’ to mechanistically describe the molecular mechanisms driving fitness of RdRp in 1) the early ‘phase I’ of pandemic before the emergence of any currently recognized VOC, 2) for ‘opportunistic’ Alpha VOC sequences that dominate the ‘phase II’ of pandemic, and 3) for ‘predatory’ Delta VOC sequences that dominate the ‘phase III’ of the pandemic (**Fig. 1A**). The ‘GPR-based fitness scores’ generated from fitness landscapes for every residue in RdRp reveal previously undetected structural ‘covariant fitness clusters’ defined by their changes in thermodynamic stability and residue connections in driving virus spread during the pandemic. These covariant fitness clusters involve structure adjustments for Zn^2+^ binding, multi-residue interactions in the N-terminal <u>Ni</u>dovirus <u>R</u>dRp <u>a</u>ssociated <u>n</u>ucleotidyl transferase domain (NiRAN) domain, and a covariant fitness cluster harboring G671S, a unique basal mutation that exists in almost all the Delta VOC sequences. We propose that GPR fitness scores defined by SCV relationships provide a computational platform to mechanistically assess the role of natural selection in the rapidly evolving viral lifecycle contributing to the pandemic.

**Figure 1.**
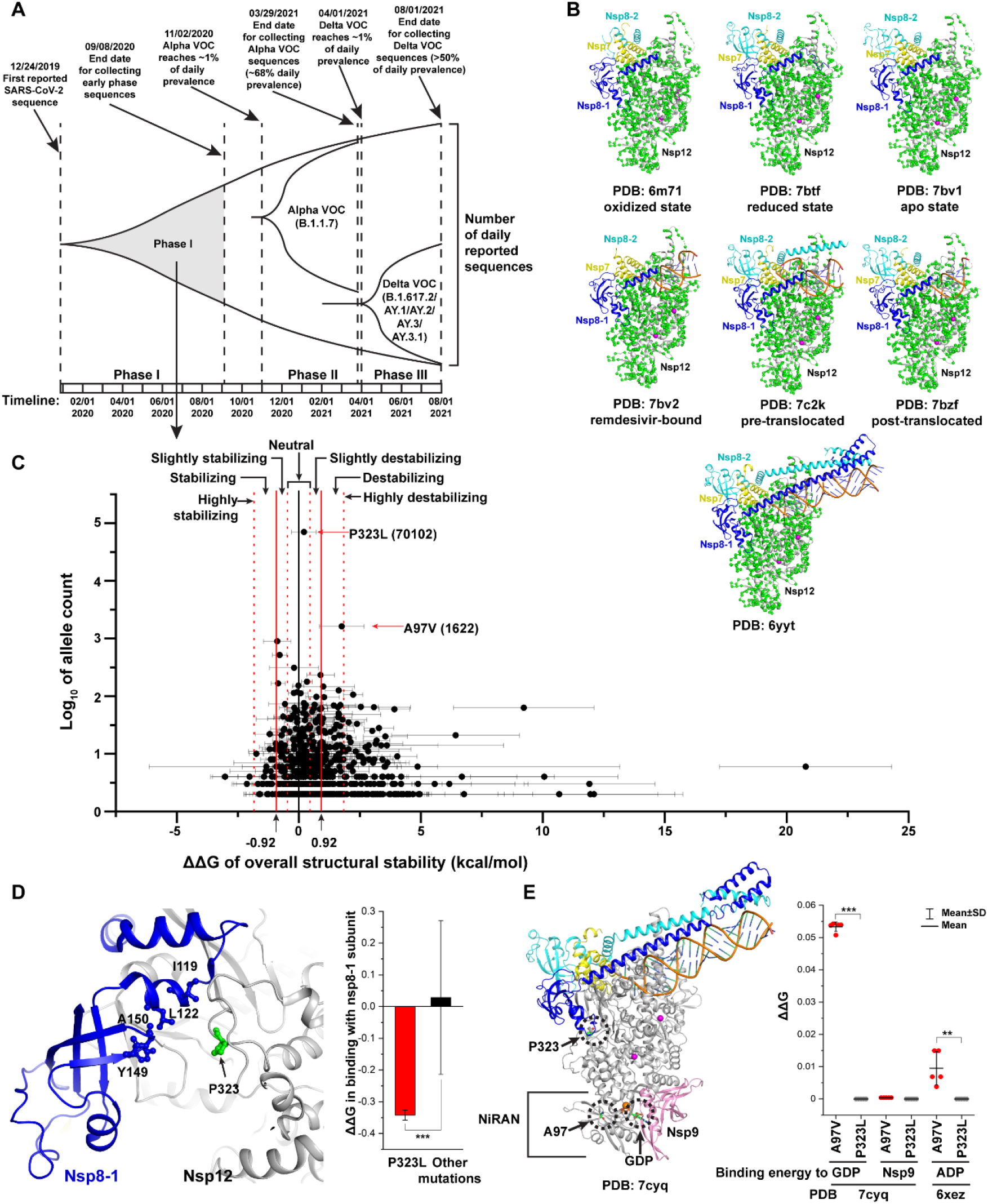
Impact of mutations on structural thermodynamic stability of nsp12. (**A**) Timeline of the early ‘phase I’ of the pandemic before the surge of Alpha VOC in the ‘phase II’ and the Delta VOC in the ‘phase III’ of the pandemic. The early phase I sequences do not contain any Alpha or Delta VOC as the end date for collecting the early phase I sequences (Sept.08, 2020) is before the first submission date for the first Alpha or Delta VOC sequences. (**B**) Structures of RdRp used for the analysis of mutation impact on structural stability. The C-alpha atoms of the amino acid residues corresponding to mutation positions of nsp12 are highlighted as green balls. Disulfide bonds in the oxidized structure (6m71) are shown as purple sticks while the bound Zn^2+^ ions in other structures are showed as purple balls. (**C**) The averaged ΔΔG of structural stability and the associated allele count for each nsp12 mutation. The mutations are classified as seven categories based on the averaged ΔΔG (**see Methods**). P323L and A97V are labeled with their allele count. (**D**) Residue P323 is located at the interface between nsp8-1 (blue) and nsp12 (gray) (left panel). The interacting residues with P323 in nsp8-1 are shown as sticks and labeled. Mutation P323L shows a significant stabilizing ΔΔG when compared with other mutations (right panel) (Student’s t-test with Welch correction for unequal variance, ***p<0.001). (**E**) Residue A97 is located in the NiRAN domain that binds to nsp9 (left panel). A97V destabilizes GDP and ADP binding when compared with P323L (right panel) (Student’s t-test, **p<0.01, ***p<0.001).

## Results

### Structural impact of RdRp mutations

To follow the evolution of nsp12 structure over the time-course of the pandemic, we first looked at 87,468 SARS-CoV-2 genome sequences from the human host in GISAID database (7, 8) spanning Dec. 24, 2019 (3) to Sept. 08, 2020, prior to emergence of individual VOC (referred to as the ‘phase I’ of the pandemic) (**Fig. 1A**). Among these collected sequences, 1,569 missense mutations were identified in nsp12, ∼55% of which only appeared in one virus sample each (10). Subsequently, we used the remaining 699 mutations present in at least two individuals (≥ two allele counts) reflecting at least one human-human transmission event for these mutations. We used the Foldx software that relies on empirically derived energy terms (55, 56) to analyze the impact of these mutations on structural thermodynamic stability (referred to henceforth as structural stability), subunit binding, and RNA binding for the RdRp complex structures in different chemical environments, such as oxidized or reduced states (PDB:6m71 and 7btf) (30), structures without or with Remdesivir (PDB:7bv1 and 7bv2) (32), structures with the stalled pre- or post- translocated states (PDB:7c2k and 7bzf) (28), and the structure of nsp12 complexed with full-length nsp7 and nsp8 (PDB:6yyt) (29) (**Fig. 1B**).

Among the 699 mutations with at least two allele counts, 695 mutations are mapped to residues that have resolved structural information and can be found along the entire structure of nsp12 (**Fig. 1B**). The impact of these mutations on the overall structural stability (ΔΔG) assessed by Foldx is highly correlated between different structures (**Fig. S1A**; Pearson correlation coefficient (Pearson’s r) = ∼0.7-0.8), demonstrating that the Foldx results are robust across different structural states. We averaged the impact on structural stability in different states for each mutation and classified the mutations using the reported accuracy of Foldx computed ΔΔG (**Fig. 1C**, see **Methods**) (56, 57). The majority of the mutations (∼57%) generated in the phase I of the pandemic have a neutral or slight impact on RdRp structural stability (-0.92 kcal/mol <ΔΔG<0.92 kcal/mol) (**Fig. 1C**). Of the remaining, a number of mutations have a significant destabilizing impact (∼19% from 0.92 to 1.84 kcal/mol) or highly destabilizing impact (∼19% > 1.84 kcal/mol), whereas others have a strong stabilizing impact (∼4% < -0.92 kcal/mol) (**Fig. 1C**).

These results indicate that mutations with significant structural impact on the stability of the RdRp complex were already circulating in the human population during the first nine months of the COVID-19 pandemic.

In the dataset reflecting at least one transmission event (**Fig. 1C**), there is no significant linear correlation between the ΔΔG value impacting structural stability and the allele count in human population (**Fig. S1B**; Pearson’s r = -0.06, p = 0.1). The most frequent mutation is P323L (allele count 70,102), occurring in ∼80% of SARS-CoV-2 genomes uploaded by early September 2020 (**Fig. 1C**). Though the overall structural stability impact of P323L is estimated as neutral (**Fig. 1C**), when compared to other mutations, P323L significantly stabilizes the binding energy between nsp12 and nsp8-1 that is critical for RdRp activity (27) (**Fig. 1D**). The second most frequent RdRp mutation is A97V present in ∼2% of SARS-CoV-2 genome samples and has a significant destabilizing impact on the overall structural stability of RdRp (**Fig. 1C**). This mutation maps to the NiRAN domain (**Fig. 1E,** residues 1 to 250). The nucleotidylation activity of the NiRAN domain is essential for viral load and propagation (58, 59), but the target remains unclear (60). Recent studies showed that the NiRAN domain acts as a guanylyltransferase to catalyze transcript capping. Its binding to nsp9 inhibits this activity (61, 62). It has also been shown that the NiRAN domain catalyzes a nucleoside monophosphate transferase reaction on the N-terminus of nsp9 (63). The A97V mutation shows no impact on the binding energy to nsp9 when compared with P323L, that is not in the NiRAN domain (**Fig. 1E**). However, it shows a significant destabilizing impact on the binding to GDP or ADP, which is critical for the nuleotidylation activity of the NiRAN domain (**Fig. 1E**). Besides A97V, ∼298 additional mutations significantly impact the stability of the RdRp structure (**Fig. 1C**).

Given the lack of general linear correlation between structural stability and allele frequency (**Fig. S1B**), we set out to use a novel GPR-based machine learning approach (48) to determine at residue-by-residue resolution whether there is a hidden pattern of spatial covariant relationships at the sequence level driving the structural fitness of RdRp during the spread of SARS-CoV-2 in the human population.

### Building a GPR fitness landscape based on structural stability of RdRp

We previously showed that a GPR-based <u>v</u>ariation <u>s</u>patial <u>p</u>rofiling (VSP) approach (48), which uses only a sparse collection of variants distributed across the human population to capture at atomic resolution the SCV relationships linking genotype variation to experimental and clinical phenotypes, can be used to map protein function on a residue-by-residue basis with defined uncertainty for the entire sequence to understand human genetic disease (48–50). To explore the SCV relationships underlying the RdRp mutations that link its sequence and structural features to spread, quantified as allele count in the human population, we used 63 mutations that have either significant stabilizing (ΔΔG>0.92 kcal/mol) or destabilizing (ΔΔG<-0.92 kcal/mol) impact with >10 allele counts reflecting transmission robustness across the human population. Among the 63 mutations, A97V has an extremely high allele count that is above two standard deviations of the distribution (**Fig. 2A**). Given that GPR is sensitive to extreme outliers (64–67), we generated two GPR models, one using the 62 mutations within the two standard deviations (**Fig. 2B-E; Fig. S2**) and another using all the 63 mutations including A97V (**Fig. S3**). To incorporate sequence information in the GPR model, we used the location of each of the mutations by its position on the primary sequence as the ***x*-axis** where the full-length sequence of nsp12 is assigned as 1.0 (**Fig. 2B**). To include the impact of both the stabilizing (4 mutations) and the destabilizing (59 mutations) mutations, we used the absolute ΔΔG as a log transformation (log_10_(1+|ΔΔG|)) (***y*-axis**) (**Fig. 2B**). A larger value indicates stronger structural impact in response to a larger change in Gibbs free energy compared to the wild-type (WT) structure. To link the sequence position (***x***-axis) and structural stability (***y***-axis) to the spread of virus using GPR, we used the allele count found in the population of each mutation as the ***z*-axis** value (**Fig. 2B,** color scale).

**Figure 2.**
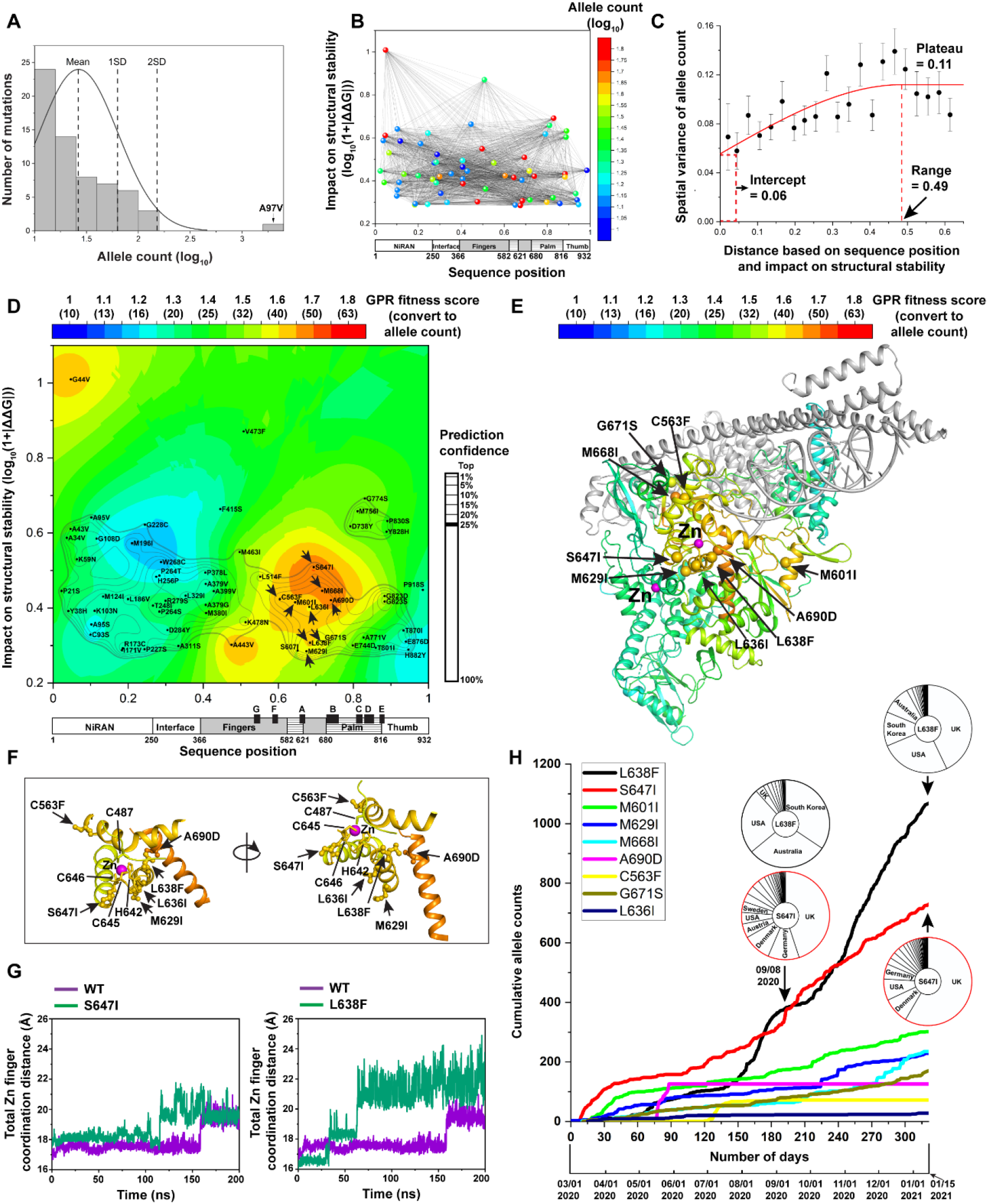
GPR fitness landscape identifies a covariant fitness cluster adjusting Zn^2^+ binding. (**A**) Distribution of the allele counts for nsp12 mutations. Mean, one standard deviation and two standard deviations of the distribution are indicated. (**B**) Nsp12 mutations are positioned by their residue positions (*x*-axis) and impact on structural stability (*y*-axis) and colored by their allele count (*z*-axis). The pairwise spatial relationships (indicated by black lines) are determined by GPR (see **Methods**). (**C**) The molecular variogram showing spatial relationships between the separation distance of paired datapoints in (**B**) and their spatial variance for the allele counts (see **Methods**). (**D**) GPR based ‘fitness landscape’ for nsp12 mutations. The predicted ‘GPR fitness score’ is indicated as color scale and the predicted spatial uncertainty (or confidence) is indicated by contour lines with the top 25% confidence indicated as bold. (**E**) The highest confidence prediction of the GPR fitness score on each residue is mapped to the RdRp structure to construct a ‘fitness structure’. (**F**) The residues where the mutations have strong GPR fitness scores around the C487-H642- C645-C646 Zn^2+^ binding motif are indicated. (**G**) Comparison of total Zn^2+^ binding coordination distance for S647I (green) (left panel) or L638F (green) (right panel) with WT (purple) found in the MD simulations of the RdRp structure. Sum of eight coordination distances (Å) to the bonded Zn^2+^ (C487-H642-C645-C646 and H295-C301-C306-C310) are plotted for WT (purple) and S647I (green) according to simulation time course. (**H**) Cumulative allele counts according to time for the mutations in the covariant fitness cluster. The country distributions of S647I and L638F at two time points (Sept.08, 2020 and Jan.15, 2021) are shown.

In our GPR-principled approach (48), a ‘molecular variogram’ (**Fig. 2C**) (48, 52) is used to analyze the relationships that link the distance between pairwise mutations in terms of their locations in primary sequence (***x***-axis) and thermodynamic structural stability (***y***-axis), to the spatial variance of the allele count (***z***-axis) (**Fig. 2B**, black lines). The molecular variogram (48, 52) represents a generalized quantitative assessment of how the nsp12 mutations correlate with each other to drive the viral spread in the context of their sequence position and impact on structural stability (see **Methods**). The variogram reveals that the spatial variance, defining the dissimilarity of the allele counts for all pairwise mutations, increases according to the mutations’ distance based on sequence positions in the nsp12 protein (***x****-axis*) and the mutations’ structural stability impact (***y***-axis) until reaching a plateau value (**Fig. 2C,** y = 0.11) at a ’range’ that spans ∼50% of the nsp12 polypeptide (**Fig. 2C, *x* =** 0.49). This result indicates that a fundamental molecular signature representing a general spatial variance relationship related to structural stability is required to drive the fitness of nsp12 mutations contributing to the spread of SARS- CoV-2.

Based on the molecular variogram generated from nsp12 mutations (**Fig. 2C**), GPR can be used to generate a structural stability fitness landscape (48) (**Fig. 2D;** see **Methods**), referred to simply as a ‘fitness landscape’ hereafter. The fitness landscape shows the predicted SCV relationships linking the structural stability impact based on ΔΔG to the allele count in human population for every residue spanning the entire polypeptide of nsp12. We refer to the predicted allele count value (***z*-axis**) in the fitness landscape for each residue in the nsp12 sequence (***x***-axis) with an assigned structural thermodynamic stability value (***y***-axis) as a ‘GPR fitness score’ (**Fig. 2D**, color scale). A higher GPR fitness score (**Fig. 2D**, yellow-orange-red) indicates an improved fitness on a residue in the human host population in response to variation when compared to variation at other residues with lower scores (**Fig. 2D**, green-cyan-blue). The GPR fitness scores in the landscape for the input mutations are strongly correlated with their actual allele counts (**Fig. S2A**, Pearson’s r = 0.73, p = 1.6 x 10^-11^), indicating that the comprehensive output fitness landscape values reliably capture the trend of the actual allele count for the sparse collection of input mutations. Comparison of the GPR model with other models such as a multivariant linear model and a decision tree based model (random forest) using leave-one-out cross-validation showed that GPR achieves the lowest root-mean-square error (**Fig. S2B-F**). Moreover, instead of generating a single value prediction, GPR assesses the probability distribution based on the spatial covariant matrix of all data points, and outputs an uncertainty (or confidence) value for each prediction (see **Methods**) that is plotted as contour lines in the GPR fitness landscape with the top 25% confidence value indicated as a bold contour line (**Fig. 2D**). The top 25% confidence value assigns value with high confidence to above 95% of the nsp12 residues. The SCV-based uncertainty value allows us to prioritize the predictions for variation found on each residue to generate residue-by-residue GPR fitness scores (see **Methods**). These residue-by-residue relationships contributing to the structural fitness of RdRp in SARS-CoV-2 provide a uniform and quantitative platform to understand the key molecular features responsible for spread in response to the environment.

### GPR fitness scores identify a covariant fitness cluster adjusting Zn^2+^ binding in RdRp

The fitness landscape reveals dramatically different GPR fitness scores within and between different domains (**Fig. 2D**). For example, we observe a major hotspot (**Fig. 2D**, yellow-orange-red) in the top 25% confidence region (**Fig. 2D**, bold contour line) that spans from the beginning of the finger domain (residue 366) to the end of palm domain (residue 815) (**Fig. 2D**, see ***x***-axis labeling). To understand the structural details of the fitness hotspot, we mapped the highest confidence prediction for each residue in the landscape onto the RdRp structure to generate a ’covariant fitness structure’ (simply referred to as a ’fitness structure’ henceforth) with the color scale representing the residue-based GPR fitness score (**Fig. 2E; Fig. S3D,** see **Methods**) (48–50). The fitness structure shows that the residues with relatively high GPR fitness scores are centered at one of the Zn^2+^ binding motifs (**Fig. 2E; Fig. S3D**). Specifically, the C563F, M629I, L636I, L638F, S647I and A690D mutations that are in the fitness hotspot map to the α-helices that are adjacent to the C487-H642-C645-C646 Zn^2+^ binding site (**Fig. 2F**). Among these mutations, L638F and S647I are at the two ends of a loop which contains H642, C645, and C646 of the Zn^2+^ binding motif (**Fig. 2F**). Molecular dynamics (MD) simulations (see **Methods**) of L638F and S647I indicate that these two mutations trigger a more rapid increase of the overall coordination distances of Zn^2+^ binding motif than WT (**Fig. 2G**). The disruption of Zn^2+^ binding by either mutation or redox-switch of the disulfide bonds for cysteine residues in the Zn^2+^ binding motif impact the overall conformational dynamics of nsp12, as well as its association with nsp8-1 subunits and RNA substrate (**Fig. S4**). Consistent with the MD simulation results, C563F in the fitness hotspot near the nsp8-1 binding site shows a significantly destabilizing impact for the binding between the nsp12 and nsp8-1 subunits (**Fig. S2G**), while A690D shows a significantly destabilizing impact for the RNA binding (**Fig. S2H**). These results indicate that SCV relationships connect the Zn^2+^ binding motif to the nsp12 binding sites for nsp8-1 and the RNA substrate (**Fig. 2E**). We posit that the residue-residue SCV relationships connecting these structural features generate a ‘covariant fitness cluster’ in the fold (**Fig. 2D** and **2E**). This covariant fitness cluster (referred as cluster 1) has significant impact on the structural stability as it consists of a cluster mutations with absolute ΔΔG values above 0.92 kcal/mol (**Fig. 2D**). Furthermore, the residues in the covariant cluster 1 have high confidence GPR fitness scores that are above the mean of input allele count (**Fig. 2D** and **E**; yellow-orange-red with z-value >1.55 (∼35 allele count)). Therefore, the SCV relationships in covariant fitness cluster 1 represent critical RdRp residue-residue structural features that evolve to augment SARS-CoV-2 fitness in the population.

Tracking the cumulative allele counts of the mutations in the covariant fitness cluster over time reveals that the allele counts of the mutations that are closest to the Zn^2+^ binding site, L638F and S647I, increase more rapidly than other mutations (**Fig. 2H**), suggesting the key roles of the Zn^2+^ binding motif in defining the evolving covariant fitness cluster 1. Interestingly, the two mutations have different time-sensitive country specific trajectories (**Fig. 2H**). For example, in early Sept. 2020, L638F was mainly found in South Korea and Australia, while S647I was mainly found in UK, Germany, and Denmark (**Fig. 2H**). These results demonstrate that a covariant fitness cluster 1 determined by the GPR fitness score represents a common feature in RdRp that SARS-CoV-2 can evolve separately in different countries reflecting the importance of SCV in evolution of the pandemic.

### N-terminal NiRAN domain fitness cluster in the Alpha VOC

The emergence of the prominent ‘opportunistic’ Alpha VOC from late 2020 ( ∼1% daily prevalence on Nov 11, 2020) to the early 2021 (∼68% daily prevalence on Mar 29, 2021) time- frame of the pandemic (**Fig. 3A**, phase II) prompted us to investigate specifically how the RdRp structure evolves in the Alpha VOC (68, 69). For this purpose, we collected 950 missense mutations in nsp12 from all the submitted Alpha VOC sequences up to Mar 29, 2021 (169,941 sequences, see **Methods**) and analyzed the structural impact of 615 mutations on nsp12 that have at least two allele counts in the Alpha VOC sequences (**Fig. 3B**). P323L, the most frequent mutation in nsp12 in the first nine months of the pandemic (**Fig. 1C**), exists in almost all the Alpha VOC sequences (**Fig. 3B**, ∼100%). Therefore, we define this mutation as a basal mutation for the Alpha VOC, highlighting the importance of stabilizing nsp12 and nsp8-1 binding (**Fig. 1D**) to promote the prevalence of the Alpha VOC. The second most frequent mutation of nsp12 in the evolution of the Alpha VOC lineage in phase II is P227L that is found in ∼5.5% of Alpha VOC sequences (**Fig. 3B**). This mutation has a slightly destabilizing structural impact (0.46 kcal/mol<ΔΔG<0.92 kcal/mol) (**Fig. 3B**) and maps to the NiRAN domain at the interface between nsp12 and nsp9 (**Fig. S5A**). When compared with A97V found in the NiRAN domain, P227L does not impact the GDP binding but significantly destabilizes the interaction to nsp9 that has been shown to be an inhibitor to the nucleotidylation activity of NiRAN domain (**Fig. S5B**).

**Figure 3.**
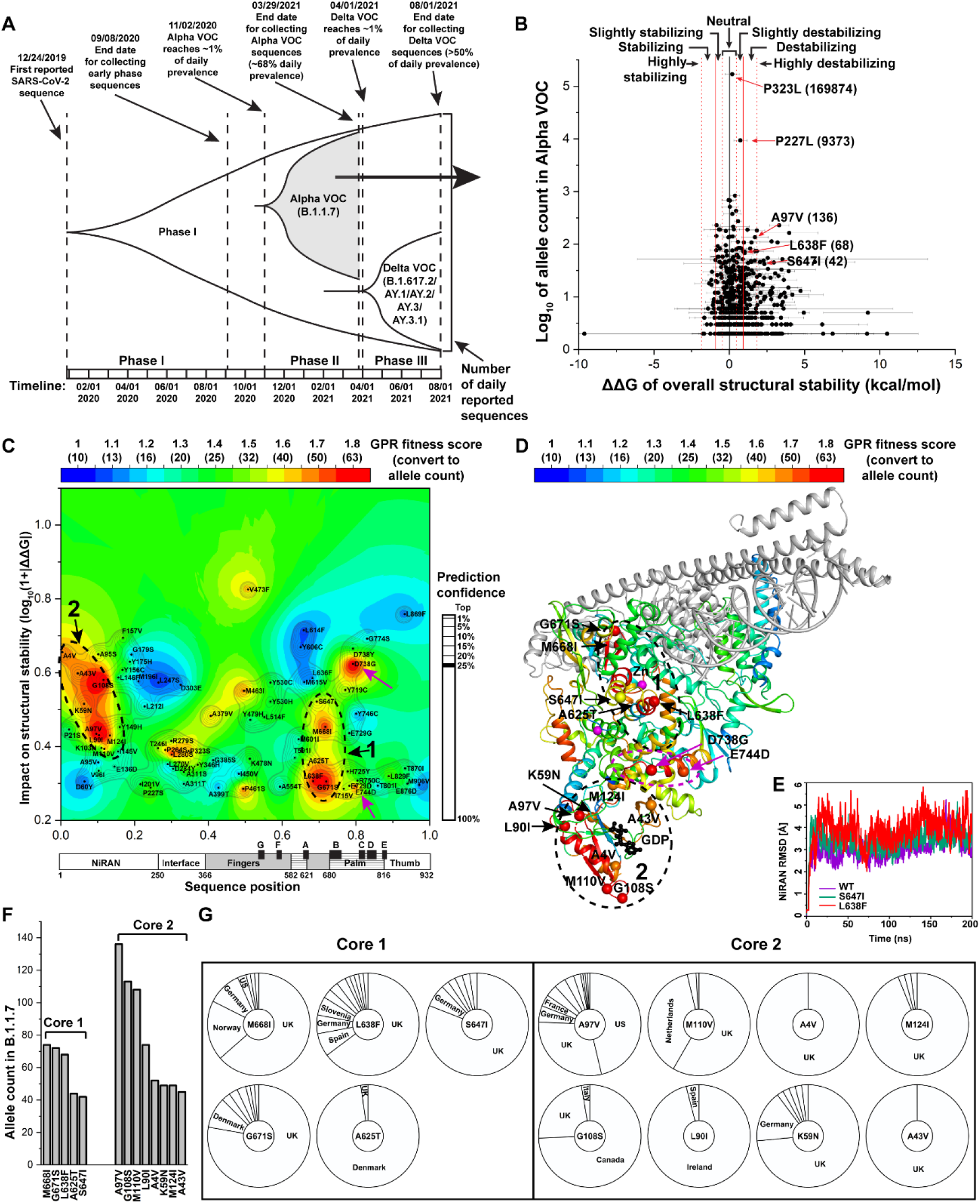
GPR fitness landscape for nsp12 mutations in the Alpha VOC. **(A)** Timeline of the surge of the Alpha VOC in the phase II of the pandemic. The date of ∼1% daily prevalence of Alpha VOC is indicated as the start date of phase II. All Alpha VOC sequences up to Mar.29, 2021 were included in the analyses. **(B)** The impact of nsp12 mutations in the Alpha VOC on the RdRp structural stability. Highlighted mutations are labeled with their allele counts. (**C**) GPR fitness landscape for the nsp12 mutations in Alpha VOC. The covariant fitness cluster around the C487- H642-C645-C646 Zn^2+^ binding motif is labeled as cluster 1 and the covariant fitness cluster in the NiRAN domain is labeled as cluster 2. (**D**) Structural mapping of the GPR fitness landscape. The mutations with high GPR fitness scores found in the covariant fitness clusters 1 and 2 are labeled. The mutations with high GPR fitness scores connecting clusters 1 and 2 are labeled and highlighted by magenta arrows. (**E**) The RMSD of NiRAN domain (residues 1-250) of WT (purple), S647I (green), and L638F (red), over 200 ns MD simulations. (**F-G**) Allele counts (**F**) and country distributions (**G**) of the mutations in covariant fitness clusters 1 and 2 in the Alpha VOC.

To understand how all of the emergent nsp12 mutations with significant impact on thermodynamic stability drive the fitness of RdRp in the Alpha VOC, we generated the GPR-based fitness landscape and the corresponding fitness structure of RdRp for the Alpha VOC (**Fig. 3C-D; Fig. S5D-H**). The covariant fitness cluster surrounding the C487-H642-C645-C646 Zn^2+^ binding motif observed in the phase I landscape (**Fig. 2D-E**) recurs in the fitness landscape and fitness structure in the Alpha VOC (**Fig. 3C-D,** cluster ‘1’). This result illustrates that the structural and functional features associated with the C487-H642-C645-C646 Zn^2+^ binding motif are key elements that SARS-CoV-2 iteratively evolves to adapt to human host environment to promote fitness feature contributing to the pandemic. Intriguingly, we observe the emergence of a new and strong covariant fitness cluster within the NiRAN domain we now refer to as covariant cluster 2 (**Fig. 3C-D**, cluster ‘2’) when compared with the phase I landscape prior to the emergence of the VOC (**Fig. 2D-E**; **Fig. S3C-D**). Given that the new covariant fitness cluster in the NiRAN domain has a significant impact on structural stability resulting in high GPR fitness scores (**Fig. 3C-D**), SCV relationships highlight an important role of the NiRAN domain in the evolution of RdRp in the Alpha VOC in the phase II of the pandemic (**Fig. 3A**).

In the nsp12 fitness structure of the Alpha VOC (**Fig. 3D**), residues in an alpha helix connecting covariant fitness clusters 1 and 2 have high predicted GPR fitness scores (**Fig. 3D**, magenta dashed circle) supported by mutations D738G and E744D (**Fig. 3C-D**, magenta arrows). These results show that there are evolving structural relationships that communicate the fitness cluster of the Zn^2+^ binding motif to the NiRAN domain. Consistent with this observation, the RMSD values of the NiRAN domain in MD simulations show that S647I (**Fig. 3E**, green line, 3.29Å) and L638F (**Fig. 3E**, red line, 3.71Å) found in Alpha VOC are larger than that of WT (**Fig. 3E**, purple line, 2.81 Å), indicating that mutations in the fitness cluster of Zn^2+^ binding motif (**Fig. 3D**, cluster 1) increase the structural plasticity of the NiRAN domain (**Fig. 3D**, cluster 2) and hence could impact the activity of the NiRAN domain to facilitate improved fitness across the population.

Though most of the mutations in covariant fitness clusters 1 and 2 reflect spread in the UK given the sequences dominating the database (**Fig. 3F-G**), additional mutations evolved in other countries. For example, A97V remains enriched in the US lineage, A625T notably predominates in Denmark, G108S evolved in Canada, and L90I is mainly restricted to Ireland (**Fig. 3G**). From a SCV perspective, these results demonstrate that unique and/or additional mutations can evolve independently in different geographical locations to facilitate the sequence based function-structure relationships dominated by the common GPR-based covariant fitness clusters.

### Delta VOC evolves a novel covariant fitness cluster 3 driven by G671S

First identified in India, the Delta VOC comprising B.1.617.2 (and AY.*) lineages spread rapidly across the globe beginning in April 2021 (**Fig. 4A,** ∼1% daily prevalence at Apr. 01, 2021) to become the dominant SARS-CoV-2 strain worldwide by August 2021 with >50% of daily sequenced SARS-CoV-2 genomes with nearly 100% prevalence in some countries (**Fig. 4A**, phase III) (70). The relative viral loads in Delta VOC cases have been shown to be higher than those in people infected with original SARS-CoV-2 strain and Alpha VOC (71–74), suggesting a faster replication cycle leading to a higher viral load in the Delta VOC.

**Figure 4.**
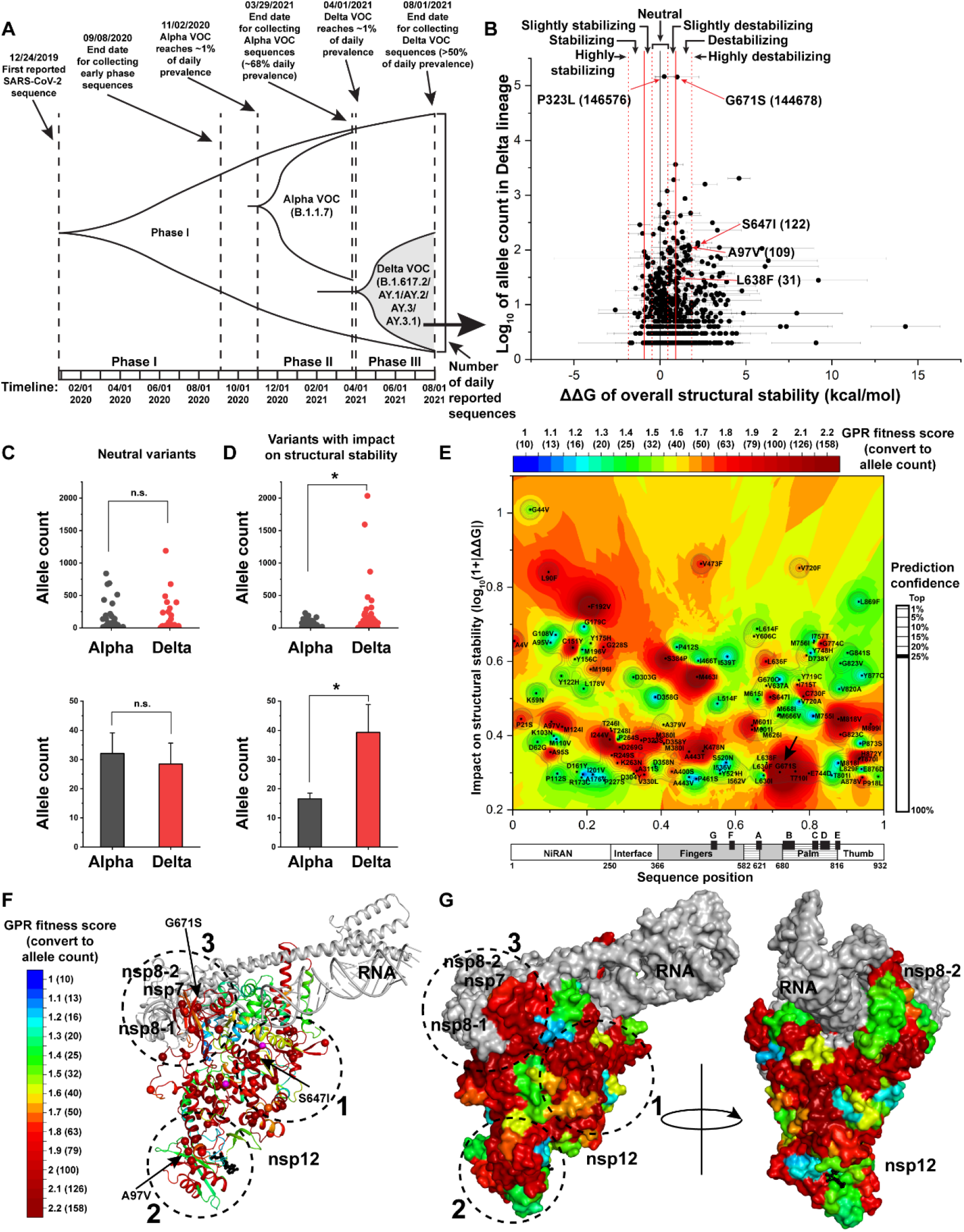
GPR fitness landscape for nsp12 mutations in the Delta VOC. (**A**) Timeline for the surge of Delta VOC. The date of ∼1% daily prevalence of the Delta VOC is indicated for the beginning of the surge. All the Delta VOC sequences (B.1.617.2/AY.1/AY.2/AY.3/AY.3.1) up to Aug.01, 2021 shown. (**B**) The impact of nsp12 mutations in the Delta VOC on the RdRp structural thermodynamic stability. (**C**) Comparison of the allele count for the neutral variants on the structural stability of the Alpha VOC and Delta VOC (Student’s t-test). The basal sequence variant P323L for both Alpha VOC and Delta VOC is not included in the analysis. (**D**) Comparison of the allele count for the variants with significant structural impact (ΔΔG>0.92 or ΔΔG<-0.92) in the Alpha VOC and Delta VOC (Student’s t-test, *p<0.05). The basal sequence variant G671S in the Delta VOC is not included in the analysis. (**E**) GPR based structural stability fitness landscape for the nsp12 mutations in the Delta VOC. G671S is highlighted by a black arrow. (**F**) Structural mapping of the fitness landscape to construct the fitness structure. S647I nearby the Zn^2+^ binding motif, A97V in the NiRAN domain and the basal mutation G671S for the Delta VOC are labeled to highlight covariant fitness clusters 1, 2, and 3, respectively. (**G**) Surface representation of the fitness structure.

To understand the evolutionary mechanism of RdRp driving Delta VOC’s ‘predatory behavior’ in the phase III pandemic, differing from the opportunistic Alpha VOC in the phase II pandemic (**Fig. 4A**), 1,073 missense nsp12 mutations were collected from 146,651 genome sequences for the Delta VOC lineages consisting of B.1.617.2, AY.1, AY.2, AY.3, and AY.3.1 up to August 01, 2021 (**Fig. 4A**) (see **Methods**) (11, 12). We analyzed the structural stability impact of 681 missense mutations on nsp12 that have at least two allele counts in the Delta VOC (**Fig. 4B**). Almost all the Delta VOC sequences (>99.9%) contain the basal P323L mutation found in the Alpha VOC. However, distinct from the Alpha VOC in which P323L is the only nsp12 basal mutation, G671S exists in >98.7% Delta VOC sequences (**Fig. 4B**), thus becoming an additional new basal mutation. The averaged ΔΔG of G671S is above 0.92 kcal/mol (**Fig. 4B**), indicating it has a significantly destabilizing effect on RdRp structure that could impact its replicative capacity. This new basal mutation in Delta VOC was found in the mutation pool of both phase I fitness landscapes (**Fig. 2D-E**) and in phase II Alpha VOC landscapes (**Fig. 3C-D**) with a high GPR fitness score at the edge of the C487-H642-C645-C646 Zn^2+^ covariant fitness cluster 1. Thus, GPR analysis indicates that the destabilizing mutation G671S has been repeatedly selected in different lineages. Integration of G671S as a basal mutation in the Delta VOC provides a unique foundational genetic framework for the evolution of RdRp activity.

In addition to the appearance of basal mutations P323L and G671S (**Fig. 4B**), the latter serving as a unique feature in nsp12 for the Delta VOC when compared with the Alpha VOC, we found that the allele count of the neutral mutations in 169,941 Alpha VOC sequences is not significantly different from that in the 146,651 Delta VOC sequences (**Fig. 4C**). In contrast, the allele count of the mutations with impact on structural stability defined by ΔΔG > 0.92 or ΔΔG <-0.92 is significantly higher than that in the Alpha VOC (**Fig. 4D**), suggesting RdRp undergoes larger structure remodeling in the Delta VOC. Consistent with this observation, the fitness landscape of the emergent Delta VOC in phase III (**Fig. 4E** and **Fig. S6A-B**) reveals a larger area with higher GPR fitness scores when compared with the phase I landscape (**Fig. 2D; Fig. S3C**) and Alpha VOC (**Fig. 3C** and **Fig. S5G**). Mapping the high confidence predictions of each residue onto the nsp12 structure reveals that the previously identified covariant fitness cluster 1 and 2 recur in the Delta VOC (**Fig. 4F-G**). Strikingly, we observe a new fitness cluster (referred to as cluster 3) around the unique basal mutation G671S in the Delta VOC (**Fig. 4F-G**). These results raise the possibility that the structural remodeling of the region around G671S contributes to the unique predatory behavior of the Delta VOC in driving the pandemic, likely through increased viral load (71–74).

### Clusters 1-3 collectively contribute to Delta VOC predatory behavior

To reveal the overall protein design of nsp12 that is responsible for the evolving predatory behavior of Delta VOC driving the pandemic, we examined the collective impact of residues with high GPR fitness scores on the different covariant fitness clusters 1-3 (**Fig. 4G**). Covariant fitness cluster 1 (**Fig. 5A-C**) includes the C487-H642-C645-C646 Zn^2+^ binding motif that we have captured from the beginning of the pandemic (**Fig. 2D-E**), in phase II Alpha VOC (**Fig. 3C-D**) and in phase III Delta VOC (**Fig. 5A**). These results validate the key importance of this region in the evolution of RdRp functional-structure design throughout the pandemic. We also observed the high GPR fitness scores for the residues that connect the Zn^2+^ binding motif to the NiRAN domain (**Fig. 5A**), illustrating evolving connectivity between these two important structural regions in both Alpha (**Fig. 3C-D**) and Delta VOC (**Fig. 5A**). Moreover, both the Alpha VOC (**Fig. 3C-D**) and Delta VOC contain a NiRAN covariant fitness cluster 2 (**Fig. 5D-F**). The interacting residues in the NiRAN domain that are evolving in the Delta VOC sequences (**Fig. 5D**) highlight the importance of this region to adapt and potentially improve function in response to the host. Covariant cluster 3 includes the basal state G671S mutation. Interestingly, A400S that interacts with G671S also shows a high fitness score (**Fig. 5G-H,** 6.2 Å between C_α_ atoms). This result suggests that acquisition of the basal sequence variant G671S in the Delta VOC sequence promotes the structural evolution of surrounding residues driving functional-structural fitness. Consistent with this observation, many residues at the interface between nsp12 and nsp8-1/nsp7 have high predicted GPR fitness scores, including the interacting residue pair, S384P and M380I (**Fig. 5G- H,** 6.0 Å between C_α_ atoms). These mutations either have lower allele counts or are absent during the transmission of the Alpha VOC sequences (**Fig. 5J-L**), resulting in significantly lower GPR fitness scores for the residues of covariant cluster 3 in the Alpha VOC when compared to the Delta VOC (**Fig. 5L**). These analyses demonstrate for the first time that the unique basal mutation G671S in the Delta VOC imposes a currently underappreciated level of evolutionary pressure driving the structural remodeling of its local environment, likely contributing to an elevated replication rate and to the increased viral load characteristic of Delta VOC (71–74).

**Figure 5.**
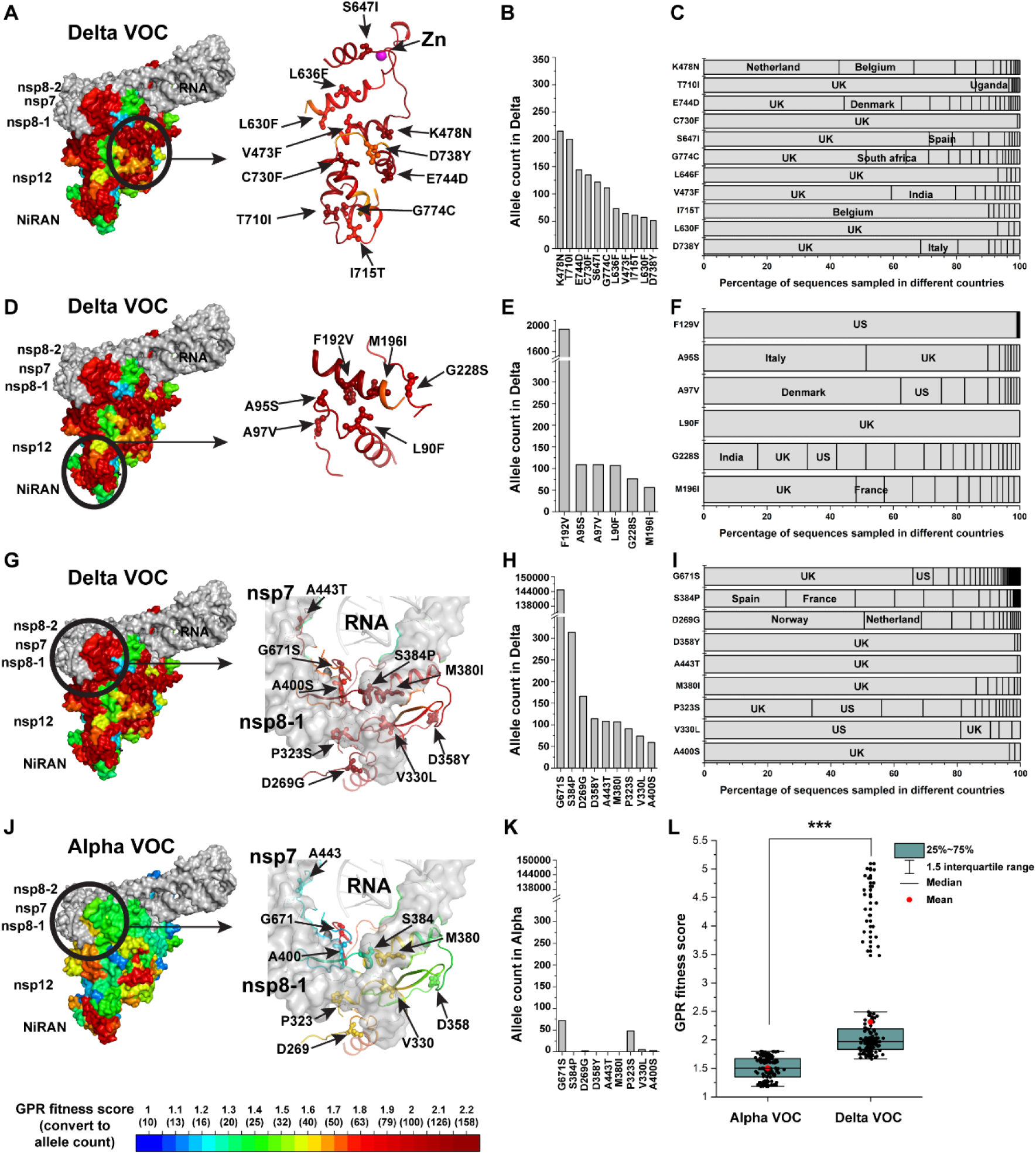
Structural covariant fitness clusters in the RdRp of the Delta VOC. (A-I) Residues with high GPR fitness scores for the Delta VOC in covariant fitness cluster 1 (**A-C**), cluster 2 (**D-F**) and cluster 3 (**G-I**) are separately illustrated. The allele count and country distribution for each mutation are presented. (**J**) GPR fitness scores in the Alpha VOC for the residues in cluster 3 are presented. (**K**) Allele counts in the Alpha VOC for the mutations in (**H**) are shown. (**L**) Comparison of the GPR fitness scores between the Alpha VOC and Delta VOC for residues in cluster 3 (Student’s t-test, ***p < 0.001).

## Discussion

Understanding the process of natural selection in response to variation responsible for the behavior of a viral lineage driving pathogen-host balance is challenging because 1) the impact of the vast majority of mutations produced during the virus life cycle appear to be neutral and 2) the changes of allele frequency due to genetic drift, biased sequence sampling, and/or a multitude of environmental events are difficult to calibrate (26, 46, 47). A commonly used method to assess the selection process measures the ratio of the non-synonymous substitutions per non-synonymous site (dN) to the synonymous substitutions per synonymous site (dS) on each codon (dN/dS) (26, 75). However, this method cannot distinguish the impact of different amino acid residues (i.e., all possible amino acid changes are treated the same as non-synonymous substitutions) (76) and hence does not directly report specific functional and coupled functional-structural features of the fold that drive fitness in the population.

Herein, we provide a method to connect mutations to inform and project the evolving sequence to functional-structural features of the fold that are important for understanding virus fitness in pathogen-host relationships. The rationale for using GPR for assessing selection is that if a functional feature (or a set of functional features) is being selected in the pathogen-host relationship, usually a cluster of mutations with similar phenotypes evolves to adjust these structural and functional features to optimize virus adaptation. The clustering effect can be quantitively framed using GPR through the principle of SCV to map with defined uncertainty critical residue-residue relationships that are shaped through evolution to drive fitness (48).

*In silico* assessment of free energy features contributing to structural stability provides one example that allows us to link the impact of viral sequence variation to the allele frequency (spread) in human population to understand fundamental functional-structural spatial relationships directing evolution on a residue-by-residue basis. For example, we observed the recurrence of C487-H642-C645-C646 Zn^2+^ binding motif as covariant fitness cluster 1 in the landscapes or structures at different time scales and across different lineages. The level of recombination events of SARS-CoV-2 during human transmission is low (77–79); for example, only 16 recombinant sequences were identified in 279,000 SARS-CoV-2 sequences from the UK dataset (79). Therefore, the recurrence of covariant fitness cluster 1 is unlikely due to the recombination with the earlier sequences; rather, the recurrence is more plausibly a result of evolutionary events that optimize SCV relationships dictating functional-structural advantage. Indeed, Zn^2+^ homeostasis in the human host has been found critical for immune responses (80), anti-viral activity (81), regulating RdRp activity (82), viral expansion (83), and the severity of disease in COVID-19 patients (83–85). A recent study demonstrated that the Zn^2+^ binding motifs in SARS-CoV-2 RdRp can also bind iron-sulfur metal cofactors instead of Zn^2+^ (86). The Fe-S bound nsp12 has a much higher specific activity than the Zn^2+^ bound form (86), suggesting that the mutations with high GPR fitness scores surrounding the Zn^2+^ binding site could selectivity evolve RdRp activity through the plasticity of these interactions. Moreover, the RNA polymerase activity of SARS-CoV-2 RdRp is nearly abolished by C487S-C645S-C646S mutations but not by C301S-C306S-310S mutations (86), consistent with our observation that covariant fitness cluster 1 is centered at C487-H642-C645- C646 Zn^2+^ binding motif and not at the other Zn^2+^ binding motif. These results suggest that the RdRp activity is allosterically managed by residue-to-residue contacts in covariant fitness cluster 1. Therefore, the efficacy of the current clinically promising RdRp nucleoside analogues targeting the active site, such as remdesivir (32, 33), favipiravir (87), and molnupiravir (41), could also be allosterically impacted by variations in covariant fitness cluster 1. More specific inhibitors could be designed to tailor the local residue-residue SCV relationships in the covariant fitness cluster 1 that are critical for the catalytic activity driving SARS-CoV-2 fitness (48, 88).

A marked difference observed in the GPR-based fitness landscape and its corresponding fitness structure for the phase I sequences compared to the Alpha VOC sequences is associated with the acquisition of destabilizing mutations in covariant fitness cluster 2 housing the NiRAN domain. The change of SCV relationships in the NiRAN domain suggest that the observed increased transmission rate of 40-50% for Alpha VOC over precursor lineages in the early phase I (69) with its unique genetic background (89) required a boost in the NiRAN activity. This interpretation is consistent with the fact that the emergence of Alpha VOC lineage during chronic infection (90) set the stage for development of covariant fitness cluster 2. Cluster 2 recurs in the Delta VOC, demonstrating that the variations in the NiRAN domain may also be important in driving the fitness of the Delta VOC (60). Besides roles in transcript capping function (61), nucleotidyltransferase activity on nsp9 (63), and/or kinase like activity (91), the NiRAN domain can bind to the N-terminal exoribonuclease (ExoN) domain of nsp14, mediating the dimerization of the replication-transcription complex (62). The potential multifunctional role(s) of the NiRAN domain that is evolving structurally during human transmission suggests that it could be a promising new drug target (91) that can be mined systematically using a GPR-based SCV platform from both *in silico* and high throughput screening (HTS) strategies to precisely target its function (48, 88).

Strikingly, the RdRp in the Delta VOC undergoes more significant structural changes when compared with the opportunistic Alpha VOC over the course of the pandemic, suggesting a rapidly evolving predatory behavior. One possibility is that plasticity in RdRp of the Delta VOC is due to the presence of the destabilizing basal mutation G671S found in >98.7% of the VOC sequences. Indeed, high GPR fitness scores are observed for the sequence and structural regions surrounding G671S that contribute to the binding interface between nsp12 and nsp8-1 subunits and that result in the new covariance fitness cluster 3. This newly evolved specialized structural pattern in the Delta VOC is consistent with an increase of viral load when compared with the population infected by the original strain identified in Wuhan and the Alpha VOC (71–74). Moreover, the structural evolution of RdRp in the Delta VOC is not restricted to the binding interface between nsp12 and nsp8-1 harboring G671S but influences multiple fitness clusters in the nsp12 structure (**Fig. 5**), suggesting a potential global structural impact of G671S. The recent Omicron VOC harboring a distinct genetic background with >30 basal mutations on the spike protein that convey novel immune evasion features (92–96) rapidly became the dominant SARS-CoV-2 strain worldwide by the beginning of 2022. Early studies show that Omicron is associated with a reduced risk of COVID-19 hospitalization when compared to Delta (97, 98). Like Delta, the Omicron VOC has P323L as the nsp12 basal mutation; unlike Delta, it is missing the G671S basal mutation. We posit that the absence of the G671S basal mutation could be one of the reasons for the attenuated pathology. In summary, our GPR approach suggests that computational and experimental surveillance of multiple SCV-based functional-structure relationships of the RdRp will be necessary to track and forecast key events that catalyze the evolving path of SARS-CoV-2 pathology.

Given the universality of our SCV-principled approach to explore the path and consequences of natural selection leading to fitness in the population in rare disease (48-50, 88, 99, 100), GPR can be applied to analyze the impact of variation in other components critical for the viral lifecycle from both basic and therapeutic perspectives (88). We focused on the fundamental thermodynamic contributions of amino acids to develop a mechanistic understanding of the SCV relationships driving the different phases of the world-wide pandemic by the Alpha and Delta VOC. Incorporation of experimental measurements developed in cell models or use of clinical data from sequenced COVID-19 patients could be a future direction to define SCV relationships impacting key physiological features contributing to pathology. Clinical input would allow us to leverage real-world, evolutionary diverse environments underpinning the design of SCV-based covariant fitness clusters to understand and therapeutically manage host-pathogen relationships, analogous to the approach we have shown for rare disease (48-50, 88, 99, 100).

## Methods

### Nsp12 mutation datasets

All nsp12 mutations in SARS-CoV-2 sequences collected from Dec.24, 2019 to Sept.08, 2020 for the early phase I study were obtained from CoV-GLUE database (http://cov-glue.cvr.gla.ac.uk/) (10) which annotates the mutation information of SARS-CoV-2 sequences from <u>g</u>lobal initiative on <u>s</u>haring <u>a</u>ll influenza <u>d</u>ata (GISAID) database (7, 8). We used the database that was last updated on Sept.08, 2020 that included 87,468 SARS-CoV-2 sequences that passed the exclusion criteria (>29,000 nucleotides, human host, covering >95% of coding region (10)) from 95526 sequences in GISAID. The collected sequences in this phase do not contain any Alpha VOC and Delta VOC sequences as the earliest submitted date for high quality Alpha VOC sequence was in Oct. 2020 and the earliest submitted date for high quality Delta VOC sequence was in Feb. 2021. Only missense mutations are considered in this study. The mutation information for the continuous tracking study from Mar.01, 2020 to Jan.15, 2021 (**Fig. 2H** and **Fig. S3G**) was also obtained from CoV-GLUE database.

The information for nsp12 mutations in the Alpha VOC comprising B.1.1.7 lineage of SARS-CoV-2 was obtained from the 2019 Novel Coronavirus Resource (2019nCoVR) at the China National Center for Bioinformatics (https://ngdc.cncb.ac.cn/ncov/?lang=en) (9, 11, 12).

2019nCoVR database integrates SARS-CoV-2 sequences from GISAID, National Center for Biotechnology Information (NCBI), National Microbiology Data Center (NMDC) and China National Center for Bioinformation (CNCB)/National Genomics Data Center (NGDC). As of the last update on Mar.29, 2021, 242,989 SARS-CoV-2 sequences were annotated as B.1.1.7 lineage by using PANGO nomenclature (101). Among these sequences, 169,941 SARS-CoV-2 sequences are complete and in high quality based on the criteria of level of unknown bases, degenerate bases, gaps, mutation density and so on (9, 11, 12). Missense mutations on nsp12 were obtained from these high quality sequences through the gff3 files provided by 2019nCoVR database.

The information for nsp12 mutations in the Delta VOC was obtained from 2019nCoVR database. 146, 651 SARS-CoV-2 sequences of B.1.617.2, AY.1, AY.2, AY.3 and AY.3.1 lineages with high sequencing quality annotated till August 01, 2021, are used to identify the missense mutations for nsp12 in Delta VOC. The nomenclature for Delta VOC used was before the changes on the nomenclature of the AY lineage series (https://www.pango.network/new-ay-lineages/).

### Calculation of structural impact of mutations with FoldX

The calculation of the free energy on RdRp structures for nsp12 mutations was performed by using FoldX 5.0 (55). Foldx has been widely used for assessing structural stability impacted by genetic mutations (102–106). The prediction results were found to be in good correlation with the *in vitro* experimental stability measurements (104, 107, 108) and have been shown to outperform the results generated by other computational stability predictors to identify disease mutations (109).

To generate a robust assessment of the structural impact of nsp12 mutations, 7 RdRp structures (PDB: 6m71, 7btf, 7bv1, 7bv2, 7c2k, 7bzf and 6yyt) were used for the analysis. The structure of 7c2k has the most complete residue information on nsp12 with only residue 908 and 909 missing. It was used as the template to fill in the missing residues in other structures by using Pymol software. The “RepairPDB” function in FoldX 5.0 was first used to correct bad torsional angles, van der Waals clashes and residues with bad energies for each structure. Then the “BuildModel” function was used to generate structural model for each mutation. This process was performed in five replicates for each mutation. The change in structural thermodynamic stability resulting from the mutation was calculated as ΔΔG (kcal/mol) = ΔG_mutant_ - ΔG_WT_ and reported as a mean ± SD over the five model pairs. The previously reported standard deviation (0.46 kcal/mol) (56) of the difference between FoldX generated ΔΔG values on structural stability and experimental measured values was used to bin the mutations into seven categories: neutral (-0.46 kcal/mol <ΔΔG< 0.46 kcal/mol), slightly stabilizing (-0.92 kcal/mol <ΔΔG< -0.46 kcal/mol), slightly destabilizing (0.46 kcal/mol <ΔΔG< 0.92 kcal/mol), stabilizing (-1.84 kcal/mol <ΔΔG< - 0.92 kcal/mol), destabilizing (0.92 kcal/mol <ΔΔG< 1.84 kcal/mol), highly stabilizing (ΔΔG< - 1.84 kcal/mol) and highly destabilizing (ΔΔG>1.84 kcal/mol)(106). The “AnalyseComplex” function was used to assess the impact of mutation on the binding energy between nsp12 and other subunits or RNA. The change in binding energy resulting from the mutation was calculated as ΔΔG (kcal/mol) = ΔG_mutant_ - ΔG_WT_ and reported as a mean ± SD over the five model pairs. The more recent RdRp structures (PDB: 7cyq and 6xez) were used to assess the mutation impact on the binding energy to ADP (6xez) or GDP (7cyq) in the NiRAN domain and the binding energy to nsp9 (7cyq).

### Molecular dynamic simulations

#### Construction of the model

The RdRp wild-type (WT) nsp12/nps8/nps7 protein complex was modeled according to the following steps. The initial protein atomic coordinates were taken from the cryo-EM PDB structure 7BTF.pdb (2.9 Å) (30). Missing internal residues in nsp12 (897–942) and residues 69-83 in nsp7 were reconstructed using the atomic coordinates from 6YYT.pdb (29) by overlapping the two structures using the Matchmaker tool in Chimera (110). The protein mutations S647I and L638F were constructed using the WT structure and mutating the residues S647 and L638, respectively, with CHARMM to replace the atomic coordinates of the side chains. While keeping all other atoms fixed, the geometry of the side chain atoms of the mutated structure was then optimized with 500 steps of steepest descent (SD) energy minimization, followed by 1000 adopted basis Newton-Raphson (ABNR). To model WT under oxidizing conditions, CHARMM was used to introduce the two disulfide bonds, between Cys301-Cys306 and between Cys487-C645 (111).

To determine an initial protonation pattern in nsp12/nsp7/nsp8, pKa values of all titratable residues were evaluated with electrostatic energy computations using in-house software karlsberg+ (112). This procedure combines continuum electrostatics with structural relaxation of hydrogen atoms and salt bridges. Assuming a pH of 7, His309, His642, and His295 were protonated on Nɛ. Under reducing conditions, the following Cys residues are assumed to be deprotonated: Cys301, Cys306, Cys487, Cys645. Under oxidizing conditions, disulfide bonds are modeled between Cys301-Cys306 and between Cys487-C645, and Cys646 and Cys310 are deprotonated. All other amino acids were protonated according to standard protonation patterns using the H-build tool from CHARMM (111).

#### Geometry optimizations and molecular dynamics

The initial geometry of each RdRp complex was solvated using the CHARMM-GUI in a water box of 90,229 explicit TIP3 water molecules, with 338 Cl^-^ and 353 Na^+^ ions to neutralized charge. The total had a total size of 290,155 atoms and was simulated in a square box of dimension 145 Å x 145 Å x 145 Å.

The solvated protein complex was next energy minimized with 10,000 steps of conjugate gradient energy minimization steps to remove any close contacts. All energy minimizations and geometry optimizations used the all-atom CHARMM36 parameter set for the protein, Cl^-^ and Na^+^ ions and the TIP3P model for water molecules (113). Van der Waals parameters for the Zn^2+^ ions were taken from the CHARMM22 parameter set, which demonstrate better agreement with experimental radial distribution functions than do the newer Zn^2+^ parameters published by Stote and Karplus (114).

After 250 ps of equilibration, the solvated protein-membrane complex was simulated with Langevin molecular dynamics (MD) at 310.15 K for 200 ns with an integration time step of 2 fs and damping coefficient of 1 ps-1. To simulate a continuous system, periodic boundary conditions were applied. Electrostatic interactions were summed with the Particle Mesh Ewald method (115) (grid spacing ∼1 Å; fftx 150, ffty 150, fftz 150). A nonbonded cutoff of 12.0 Å was used, and Heuristic testing was performed at each energy call to evaluate whether the non-bonded pair list should be updated.

### Building fitness landscapes and fitness structures using GPR based VSP

The VSP analysis of nsp12 mutations was performed as previously described(48–50) using gstat package (V2.0) in R. VSP is built on Gaussian process regression (GPR) based machine learning. A special form of GPR machine learning that has been developed in geostatistics, Ordinary Kriging (52), is used to model the spatial dependency as a variogram to interpolate the unmeasured value to construct the fitness landscape.

#### Variogram analysis

Nsp12 mutations were positioned by their sequence positions in the polypeptide chain on the ‘x’ axis coordinate and their impact on structural stability on the ‘y’ axis coordinate to the allele count along the ‘z’ axis coordinate. Suppose the i^th^ (or j^th^) observation in a dataset consists of a value z_i_ (or z_j_) at coordinates x_i_ (or x_j_) and y_i_ (or y_j_). The distance h between the i^th^ and j^th^ observation is calculated by:

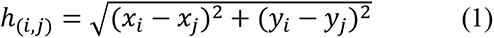

The *γ*(*h*)*-*variance for a given distance (*h*) is defined by:

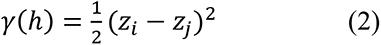

where (*h*)-variance is the semivariance (i.e., the degree of dissimilarity) of the *z* value between the two observations, which is also the whole variance of *z* value for one observation at the given separation distance *h,* referred to as spatial variance here. The distance (*h*) and spatial variance (*γ*(*h*)) for all the data pairs are generated by the equations (1) and (2). Then, the average values of spatial variance for each distance interval are calculated to plot the averaged spatial variance versus distance. The fitting of variograms were determined using GS+ Version 10 (Gamma Design Software) by both minimizing the residual sum of squares (RSS) and maximizing the leave-one- out cross-validation result (see below). Spherical and Exponential variogram models were used in this study.

The formular of the spherical model is:

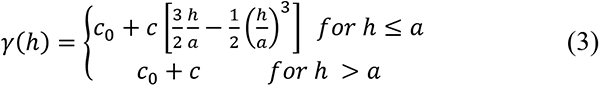

where *c*_0_ is the y-intercept of variogram (or the nugget constant); *c*_0_ + *c* is the plateau (or the sill) of variogram; *a* is the effective range.

The formular of the Exponential model is:

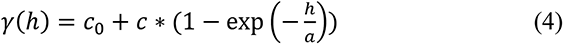

where *c*_0_ is the y-intercept of variogram (or the nugget constant), *c*_0_ + *c* is the plateau (or the sill) of variogram, *a* is the range (effective range is 3*a*).

The molecular variogram defines quantitatively the correlation between the spatial variance of z changes and the separation distance defined by the x and y coordinates based on known mutations. The distance where the model curve first flattens out is known as the range. Locations separated by distances closer than the range are spatially correlated, whereas locations farther apart than the range are not. The variogram enables us to compute the spatial covariance (SCV) matrices for any possible separation vector. The SCV at the distance (h) is calculated by C(*h*)= C(0) − *γ*(*h*), where C(0) is the covariance at zero distance representing the global variance of the data points under consideration (i.e., the plateau of the variogram).

#### Assessing the uncertainty

GPR based Kriging aims to generate the prediction that has minimized estimation error, i.e., error variance, which is generated according to the expression:

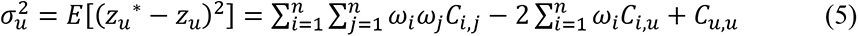

where z*_u_ is the prediction value while *z_u_* is the true but unknown value, *C_i,j_* and *C_i,u_* are SCV between data points i and j, and data points i and u, respectively, and *C_u,u_* is the SCV within location *u*. *ω_i_* is the weight for data point i. The SCV is obtained from the above molecular variogram analysis and the weight (*ω_i_*) solved from equation (5) is used for following prediction. To ensure an unbiased result, the sum of weight is set as one:

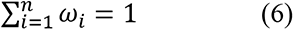

Equations (5) and (6) not only solved the set of weights associated with input observations, but also provide the minimized ‘molecular variance’ at location u which can be expressed as:

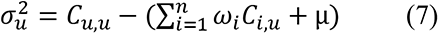

where *C_u,u_* is the SCV within location *u*, *ω_i_* is the weight for data point i, and *C_i,u_* are SCV between data points i and u. μ is the Lagrange Parameter that is used to convert the constrained minimization problem in Equation (5) into an unconstrained one. The resulting minimized molecular variance assessing the prediction uncertainty presents the confidence level of the prediction.

#### The matrix notation

The minimization of GPR-based Kriging variance (equation (5)) with the constraint that the sum of the weights is 1 (equation (6)) can be written in matrix form as

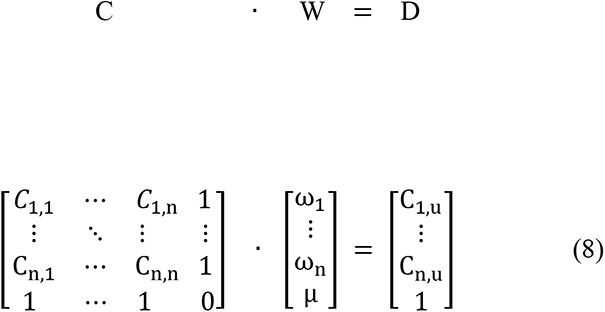

where ‘C’ is the covariance matrix of the known data points. ‘W’ is the set of weights assigned to the known data points for generating the predicted phenotype landscape. ‘μ’ is the Lagrange multiplier to convert a constrained minimization problem into an unconstrained one. ‘D’ is the covariance matrix between known data points to the unknown data points. Since ‘W’ is the value we want to solve to generate the fitness landscape, this equation can be also written as

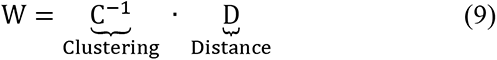

Where ‘C^-1^’ is the inverse form of the ‘C’ matrix.

As a more intuitive explanation of the GPR-based Kriging matrix notation, herein we simply refer to the VSP matrix that generates the fitness landscape (‘W’) to be based on the two important computational features used for predicting the unknown fitness values from the known-(1) the clustering (i.e., clustered sequence values with similar fitness properties ‘C^-1^’) and (2) the distance constraints (D). Here, ‘C^-1^’ represents the clustering information of the known data points while ‘D’ represents predicted statistical distance between known data points to unknown data points.

#### Generating the prediction

With the solved weights W, we can calculate the prediction of all unknown values to generate the complete fitness landscape by the equation:

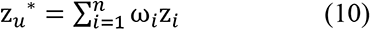

where z*_u_ is the prediction value for the unknown data point *u*, *ω_i_* is the weight for the known data point, and *z_i_* is the measured value for data point i.

#### Prediction validation and input data requirements

Leave-one-out cross-validation (LOOCV) is used because of small sample size modeling (116). In the LOOCV, we remove each data point, one at a time and use the rest of the data points to predict the missing value. We repeat the prediction for all data points and compare the prediction results to the measured value to generate the Pearson’s r-value and its associated p-value (ANOVA test performed in Originpro version 2020b (OriginLab)). Sampling data input required for reliable GPR-based Kriging or VSP prediction not only depends on the sample size (number of boreholes/locations/(variant equivalents for studies herein)) but also depends on the spatial distribution of the samples. In the previous study (48), we have showed that the VSP prediction accuracy is stable until the number of training data points drops below ∼50, consistent with the empirical rule of ∼50 data points and above recommended in geostatistical studies.

#### Mapping fitness landscape onto structure

Fitness landscapes predicted based on a sparse collection of mutations contain experimental information that comprises the full range of values on y- and z-axis for the entire polypeptide sequence (x-axis). To map the fitness predictions onto structure, we assign the prediction value with highest confidence to each residue to generate a fitness structure displaying at atomic resolution that illustrates values on all the residues interpolated in the fitness landscape from the sparse collection mutations. For the structural mapping of the fitness landscape of the Alpha VOC and Delta VOC, there are cases where multiple different high confidence values are located on a single residue. For these residues, we select the prediction with highest fitness score that are within top 1% confident region. PDB:6yyt is used for fitness structure mapping. The structural presentations were produced by the software of PyMOL.

## Data and materials availability

All the input source data, R-code scripts and output files are shared through the link: https://www.dropbox.com/sh/s2j7vrw5pa6ky60/AAAuZUiGYj0rP6EIikWOjm75a?dl=0

## Acknowledgement

We thank Daniel Shak and Dr. Salvatore Loguercio for the help on working with the mutation information from 2019nCoVR database from CNCB. We gratefully acknowledge all the authors from the originating laboratories responsible for obtaining the SARS-CoV-2 samples and the laboratories where SARS-CoV-2 genetic sequence data were generated and shared via the GISAID initiative, NCBI or CNCB, on which this research is based. Support was provided by NIH grants DK051870; HL141810; HL095524; AG070209; AG049665.

## Author contribution

C.W., N.E.-M and W.E.B designed the study. C.W. collected the mutation information, performed Foldx and GPR analysis. N.E.-M performed the MD simulation. C.W., N.E.M., and W.E.B. wrote the manuscript.

## Supplementary figures and legends

**Figure S1.**
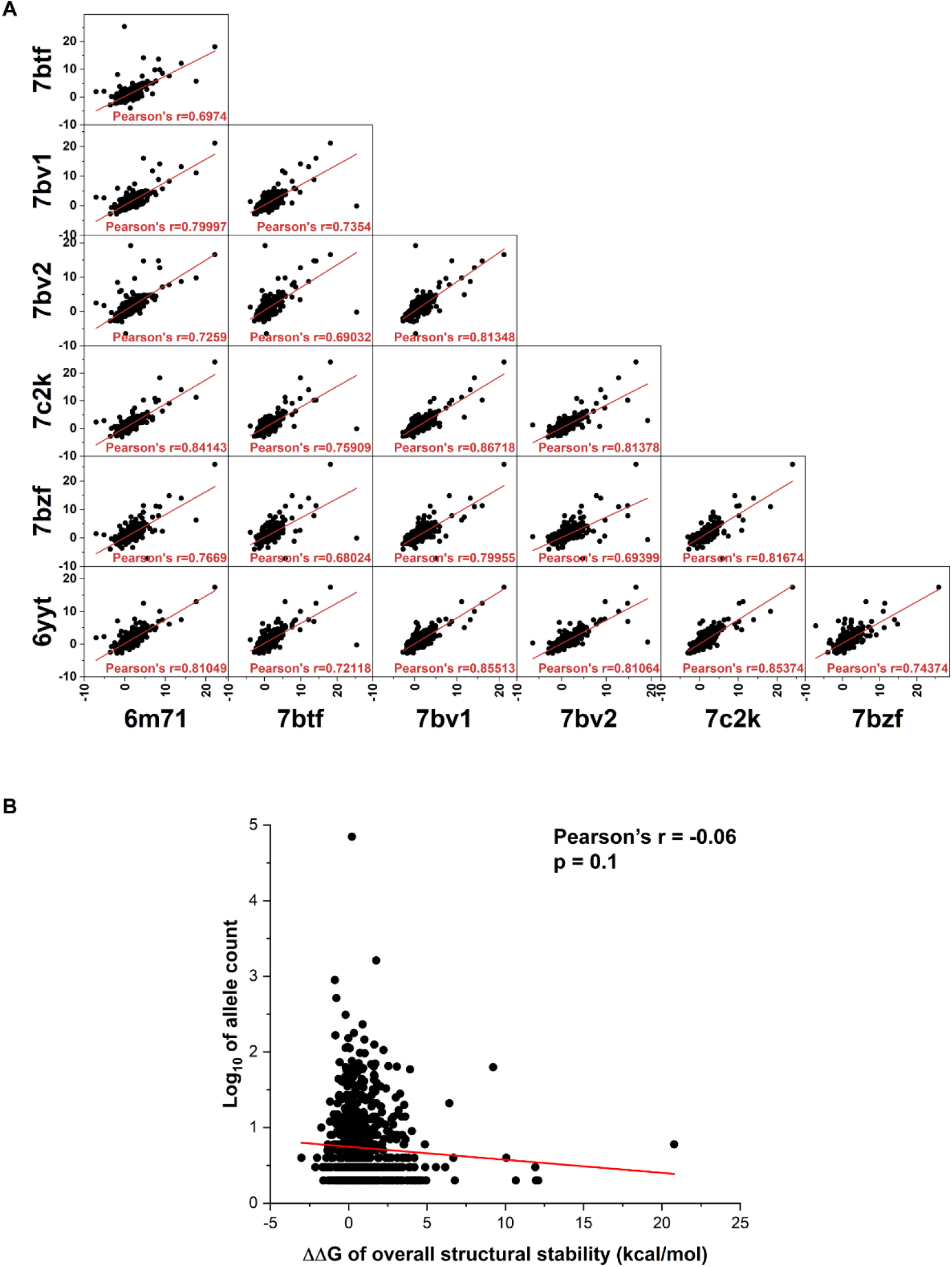
Correlation of the computed ΔΔG (kcal/mol) for mutations in nsp12 between different structural states. (**A**) The PDB ID for each structure is labeled and Pearson’s r of each correlation is indicated. (**B**) Linear correlation between averaged ΔΔG for nsp12 mutations and their allele counts. Pearson’s r and the p-value with null hypothesis of r = 0 (ANOVA test) are indicated.

**Figure S2.**
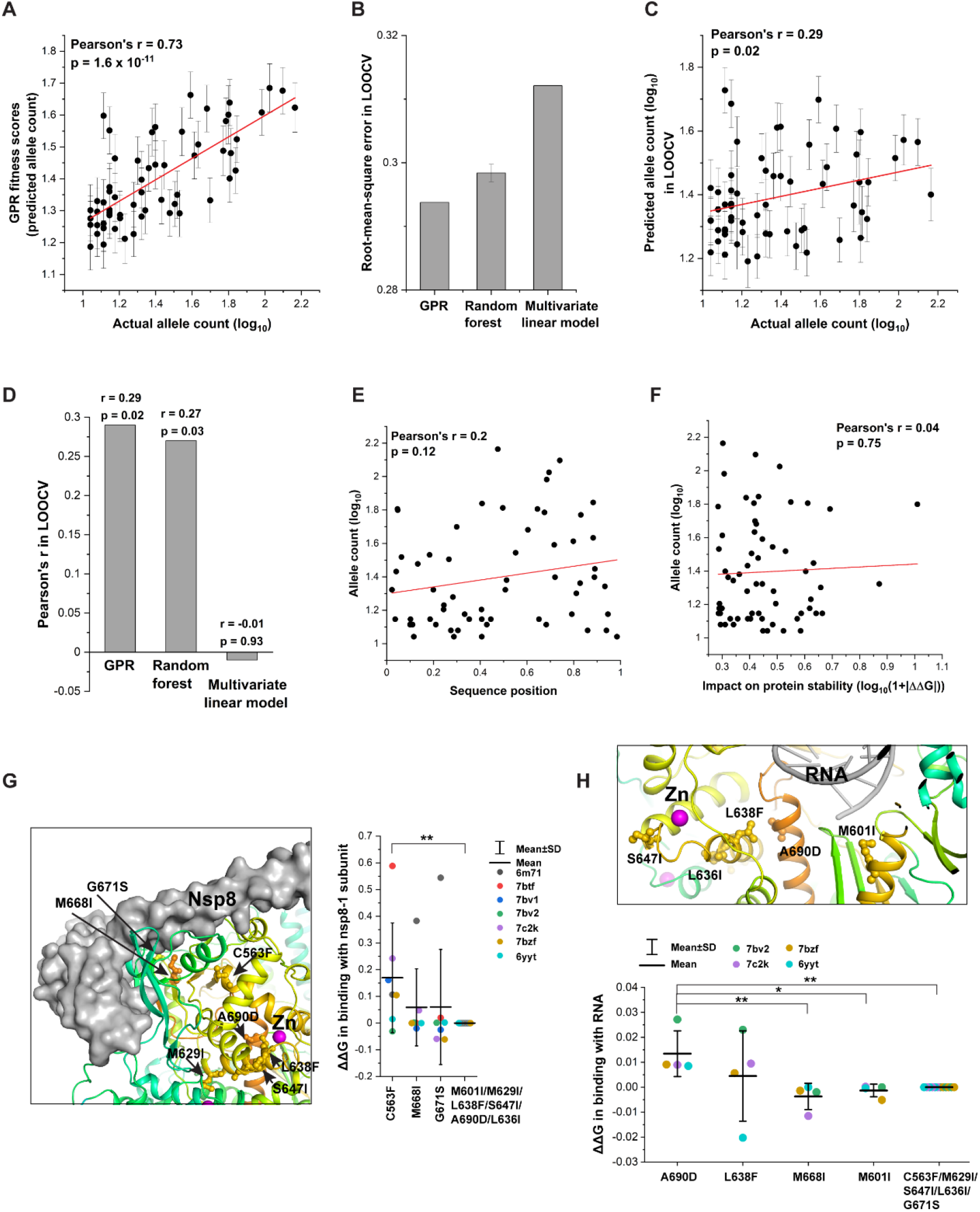
Leave-one-out cross-validation (LOOCV) of the GPR fitness landscape. **(A)** Correlation between the GPR fitness scores (i.e., predicted allele counts) in the fitness landscape for the input mutations and their actual allele counts. Pearson’s r and the p-value with null hypothesis of r = 0 (ANOVA test) are indicated. Given that there is certain spatial variance at close to “0” distance, as indicated in the variogram (Fig. 2C, y-intercept = 0.06), the GPR fitness scores in the landscape for the input mutations are not the same as their actual allele counts, but they are strongly correlated (Pearson’s r = 0.73, p = 1.6 x 10^-11^). (**B**) Root-mean-square error of GPR model, random forest model and multivariate linear model for LOOCV. (**C**) Correlation between the actual allele count and the predicted allele count in the LOOCV for the GPR model. Pearson’s r and the p-value with null hypothesis of r = 0 (ANOVA test) are indicated. (**D**) Comparisons of the correlation between the actual and predicted allele count in the LOOCV for GPR model, random forest model and multivariate linear model. (**E**) Correlation between the mutation sequence position and the allele count of mutation. (**F**) Correlation between the impact of mutation on structural stability and the allele count of mutation. While the predictions generated by both GPR and random forest achieve weak but significant correlations with the actual allele counts in LOOCV (**D**, Pearson’s r = 0.29, p = 0.02 (GPR); Pearson’s r = 0.27, p = 0.03 (random forest)), the multivariant linear model does not achieve a significant prediction (**D;** Pearson’s r = -0.01, p = 0.93). Furthermore, there is no significant direct linear correlation either between mutation position and allele count (**E;** Pearson’s =0.2, p = 0.12) or between structural stability and allele count (**F;** Pearson’s r = 0.04, p = 0.75). These results demonstrate that SCV principled GPR modeling can capture the non-linear pattern in the dataset to link sequence position information of each of the mutations with the computed structural stability to the allele count. (**G**) Impact of the mutations in the ‘covariant fitness cluster’ on the binding energy between nsp8-1 and nsp12. C563F shows a significant destabilizing impact when compared with other mutations in the covariant fitness cluster (One-way ANOVA Tukey test, **p<0.01). (**H**) Impact of the mutations in the covariant fitness cluster on the binding energy to RNA. A690D shows a significant destabilizing impact when compared with other mutations (One-way ANOVA Tukey test, **p<0.01, *p<0.05).

**Figure S3.**
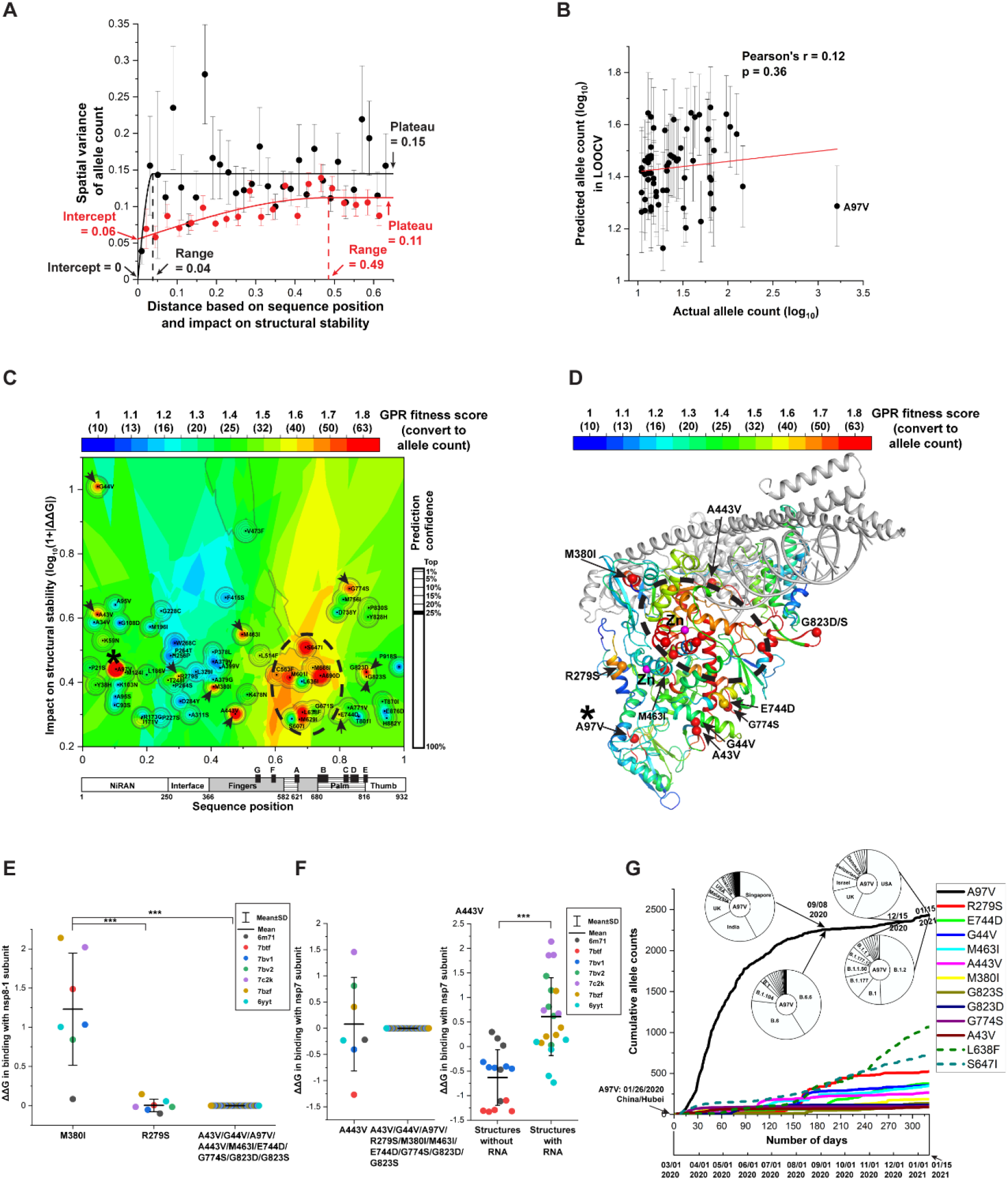
GPR model including A97V. (**A**) Molecular variograms with (black curve) or without (red curve) A97V. Including A97V in the molecular variogram analysis results in a dramatic decrease on both the correlation distance range (from 0.49 to 0.04) and y-intercept (from 0.06 to 0), and a significant increase on the plateau value representing the global variance of allele count (from 0.11 to 0.15) (compare red curve with black curve). This result suggests that A97V with an exceptionally high allele count compacts long-range SCV relationships found in response to the other mutations to very short-range SCV relationships. (**B**) Correlation between the actual allele count and the predicted allele count in the LOOCV for the GPR model including A97V. Pearson’s r and the p value with null hypothesis of r = 0 (ANOVA test) are indicated. In the LOOCV result, the allele count of A97V is underestimated by the GPR (**B**), indicating that the allele count of A97V is dramatically higher than the surrounding variants. This observation suggests that there is not much evolution going on for the A97V interacting or surrounding residues, indicating potential random events contributing to the spread of A97V in the early phase I of the pandemic. (**C-D**) GPR fitness landscape (**C**) and fitness structure (**D**) for the GPR model including A97V. Mutations with high GPR fitness scores that are not observed in the A97V excluded model (Fig. 2D**-E**) are highlighted by arrows and A97V is highlighted by *. Consistent with the fitness landscape and fitness structure without A97V (Fig. 2D**-E**), the fitness landscape and fitness structure including A97V reveals high GPR fitness scores for the covariant fitness cluster around the Zn^2+^ binding motif (**C-D,** dash circle). However, because of the short range in the variogram (**A**, black curve), the GPR fitness landscape with A97V shows additional mutations with high GPR fitness scores (**C-D**, arrows) that were not observed in the GPR model without A97V (Fig. 2D**-E**). These results reveal additional structural features that could contribute to the fitness of RdRp, for example: (**E**) M380I shows significant destabilizing impact on the binding energy with nsp8-1 (One-way ANOVA Tukey test, ***p<0.001). (**F**) A443V shows stabilizing impact on the binding energy with nsp7 for structures without RNA while shows destabilizing impact for structures with RNA (Student’s t-test, ***p<0.001). (**G**) Cumulative allele counts for mutations according to time. The lineages and country distributions for A97V at Sept. 08, 2020, and between the period from Dec. 15, 2020, to Jan. 15, 2021, are indicated. A97V was first identified in the original epicenter of COVID-19, Hubei, China on Jan. 16, 2020. Then it rapidly spread to Singapore and India as shown at the time point of Sep. 08, 2020. There is a potential founder effect contributing to this phase of rapid spread of A97V given its introduction into new geographical locations (e.g., Singapore and India) at the beginning of the pandemic. While its cumulative allele count reached a plateau in early Sept 2020, after the middle of Dec. 2020, we observe an increase of its allele counts in the US and UK that can be found in multiple different SARS-CoV-2 lineages. In the early phase I of the pandemic, A97V is dominant in B.6.6 and B.6 lineages, but between Dec. 2020 and Jan. 2021, it arises in the B.1.2 and B.1 lineages. Importantly, A97V recurs in the Alpha VOC (B.1.1.7) that harbors an unique genomic signature of 17 defining mutations (101). Alpha VOC rapidly became one of the dominant SARS-CoV-2 lineages driving the pandemic in late 2020 and early 2021. The recurrence of this significant destabilizing mutation in multiple lineages indicates a functional and structural selection at this site.

**Figure S4.**
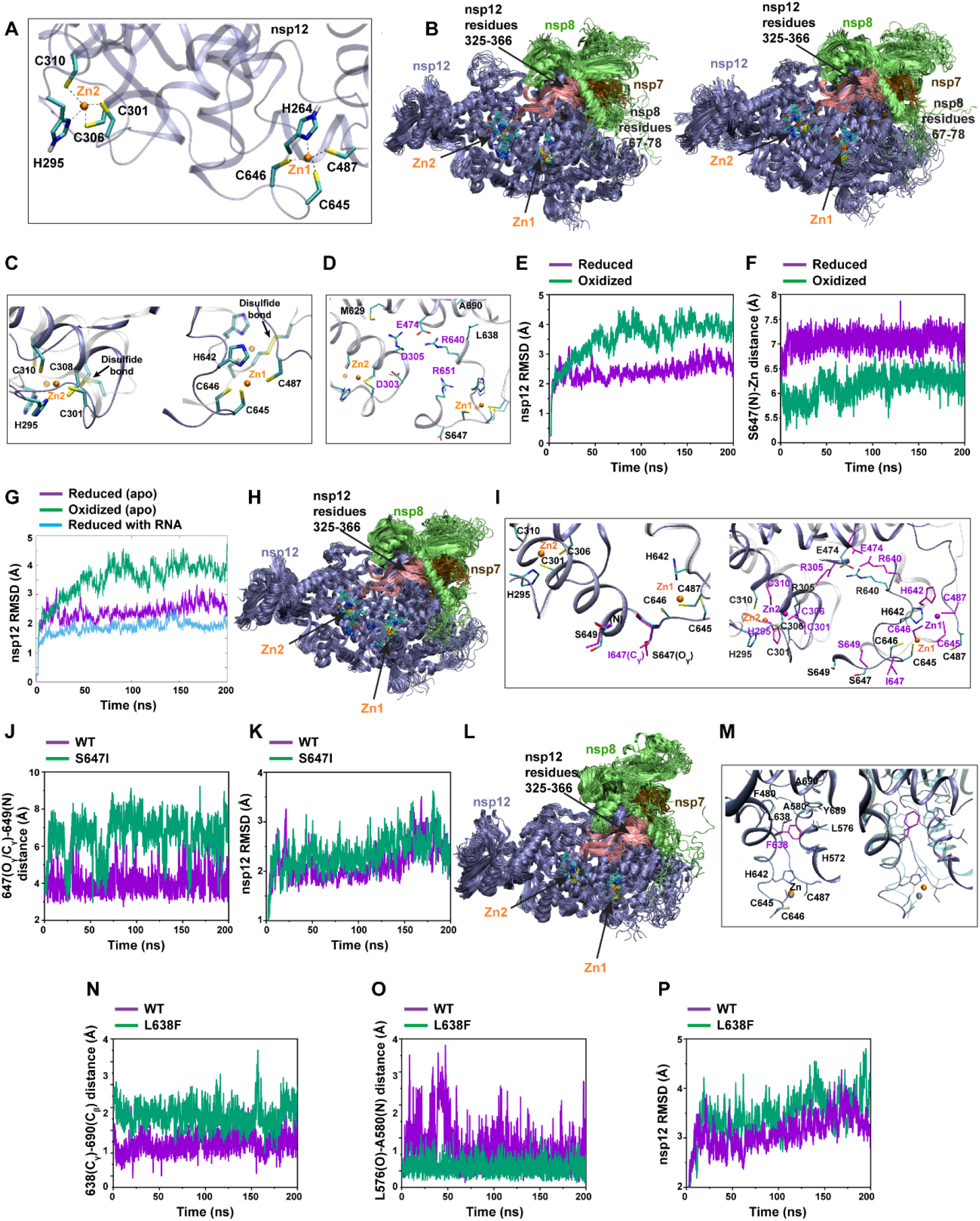
MD simulations reveal the impact of Zn^2+^ binding motifs on RdRp structures. **(A)** Zn^2+^ binding motifs up-close for WT. The two Zn^2+^ (Zn^2+^ atoms shown in orange) binding motifs in nsp12 are comprised of residues C487-H642-C645-C646, coordinated to Zn1, and H295-C301-C306-C310, coordinated to Zn2, respectively. Since C487-C645 and C301-C306 in the two Zn^2+^ binding motifs can form disulfide bonds under oxidizing conditions (30), we first performed the simulations of WT RdRp under either reducing or oxidizing conditions to represent a redox-switch that regulates the Zn^2+^ binding. Under reducing conditions, all cysteine residues are deprotonated, and the histidine side chains are protonated at the Nɛ position. Under oxidizing conditions, disulfide bonds are introduced between Cys301-Cys306 and between Cys487-C645, while Cys646 and Cys310 remain deprotonated. (**B**) Time lapse (20 snapshots taken over 200 ns) depictions of WT RdRp in the reduced state (left) and oxidized state (right). We observed the nsp12 sheets comprised of residues 325-366 (**B**, pink β-strands) in contact with nsp8 are separated and less compact in the oxidized state than that in the reduced state. Moreover, residues 67-78 of nsp8-1 (**B**, green subunit) which compose a truncated version of the long “poles” required to guide the replicating and exiting RNA (29), display different dynamic behaviors in oxidized versus reduced nsp12, showing that the chemistry of the Zn^2+^ binding motif in nsp12 directly affects the dynamics of these residues in nsp8-1. (**C**) Snapshot of WT reduced (opaque) versus WT oxidized (transparent) at 200 ns. The Zn^2+^ binding coordination is preserved over the course of the MD simulation of the reduce structure (**C**, opaque presentation, Cys301(Sγ)-Zn^2+^ ∼2.6 Å, Cys306 (Sγ)- Zn^2+^ ∼2.8 Å, Cys645(Sγ)-Zn^2+^ ∼2.5 Å and Cys487(Sγ)-Zn^2+^ ∼2.5 Å). However, in the oxidized state, the disulfide bridges Cys301-Cys306 and Cys487-Cys645 disrupt the coordinating Zn^2+^ binding motif, likely due to the lack of negative charge, resulting in the bridging disulfide cysteine residues pulling away from Zn^2+^ (Cys301(Sγ)-Zn^2+^ ∼4.6 Å, Cys306(Sγ)-Zn^2+^ ∼4.1 Å, Cys645(Sγ)- Zn^2+^ ∼5.8 Å and Cys487(Sγ)-Zn^2+^ ∼6.4 Å) (**C**, transparent representation). (**D**) Charged side chains (labeled in purple) connecting the two Zn^2+^ binding motifs. In both redox states, the charged sidechains located between the two Zn^2+^ binding motifs (Asp303, Arg305, Glu474, Asp640, Arg651) interact dynamically (**D**, residues labeled in purple), suggesting that the two Zn^2+^ binding motifs may communicate with each other through the helices based on these electrostatic interactions. (**E**) The root-mean-square deviation (RMSD (Å)) of nsp12 in WT RdRp is compared for reducing conditions (purple) versus oxidizing conditions (green) over 200 ns MD simulation. In the apo form (no RNA substrate), WT RdRp in the reduced state demonstrates overall more conserved behavior with smaller RMSD of atomic positions than WT in the oxidized state, and the side chains and backbone are more tightly clustered in the reduced state. (**F**) S647(N)-Zn^2+^ distance (Å) for reduced WT (purple) versus oxidized WT (green). Residue Ser647 is located closer to Zn^2+^ (Zn^2+^-S647 N distance ∼6 Å) in the oxidized state (**F**, green line), but under reducing conditions, the peptide backbone shifts away (Zn^2+^-S647 N distance ∼7 Å) (**F**, purple line) suggesting that the nsp12 environment near the 647 site is critical in altering nsp12 structural behavior. (**G**) Comparison of dynamic behavior of reduced WT (purple), oxidized WT (green), and reduced WT (cyan) with RNA. Simulation of WT RdRp (under reduced conditions) with RNA substrate shows more compact protein structural relationships (cyan curve), which is distinct from the more dynamic structures of the oxidized conditions (green curve). (**H**) Time lapse (20 snapshots taken over 200 ns) depictions of S647I mutation of RdRp under reducing conditions. (**I**) Comparison of Zn^2+^ binding motifs for at t=0ns (left) and t=200ns (right) for WT (opaque) and S647I (transparent). The residues in S647I in t=200ns are labeled in purple. MD simulations of the S647I mutation show that substitution of Ser647 with Ile disrupts the hydrogen bonding interaction of WT Ser647- Oγ with the amide nitrogen of Ser649. (**J**) Comparison of 647(Oγ/Cγ)-649(N) distance (Å) for WT (purple) and S647I (green). S647I mutation has larger 647(Cγ)-649(N) distance. (**K**) The RMSD of WT nsp12 (purple) and S647I nsp12 are compared over 200 ns MD simulation. Local nsp12 backbone, particularly for residues 400-700, is less constrained in S647I and can explore a larger conformational space while retaining its native function (average RMSD 2.45Å for S647I (green) vs. 2.19Å for WT (purple)). (**L**) Time lapse (20 snapshots taken over 200 ns) depictions of L638F mutation of RdRp under reducing conditions. (**M**) Comparison of Zn^2+^ binding motifs for at t=0ns (left) and t=200ns (right) for WT (opaque) and L638F (transparent). L638F, resulting in the bulky Phe638 sidechain, causes the nearby nsp12 backbone to shift relative to WT Leu638. (**N**) Comparison of L638/F638(Cγ)-Ala690(Cβ) distance (Å) for WT (purple) and L638F (green). L638F has an increase in the distance between Phe638(C_γ_) and Ala690(C_β_) that is on the nearby helix. (**O**) Comparison of Leu576(O)-Ala580(N) distance (Å) for WT (purple) and L638F (green). L638F has a decrease of the distance between Leu576(O) and Ala580(N) that are on the nearby helix. (**P**) The RMSD of WT nsp12 (purple) and L638F nsp12 are compared over 200 ns MD simulation. The RMSD (Å) values of nsp12 backbone atoms of residues 350-400 reflect an increase in conformational flexibility for L638F compared to WT. Taken together, the mutations in the Zn^2+^ covariant fitness cluster preserves the overall native structural integrity of the nsp subunits while simultaneously increasing conformational freedom to improve RdRp fitness as indicated by the GPR fitness score, providing a mechanistic interpretation to evolutionary role in the pandemic (Fig. 2D and 2E).

**Figure S5.**
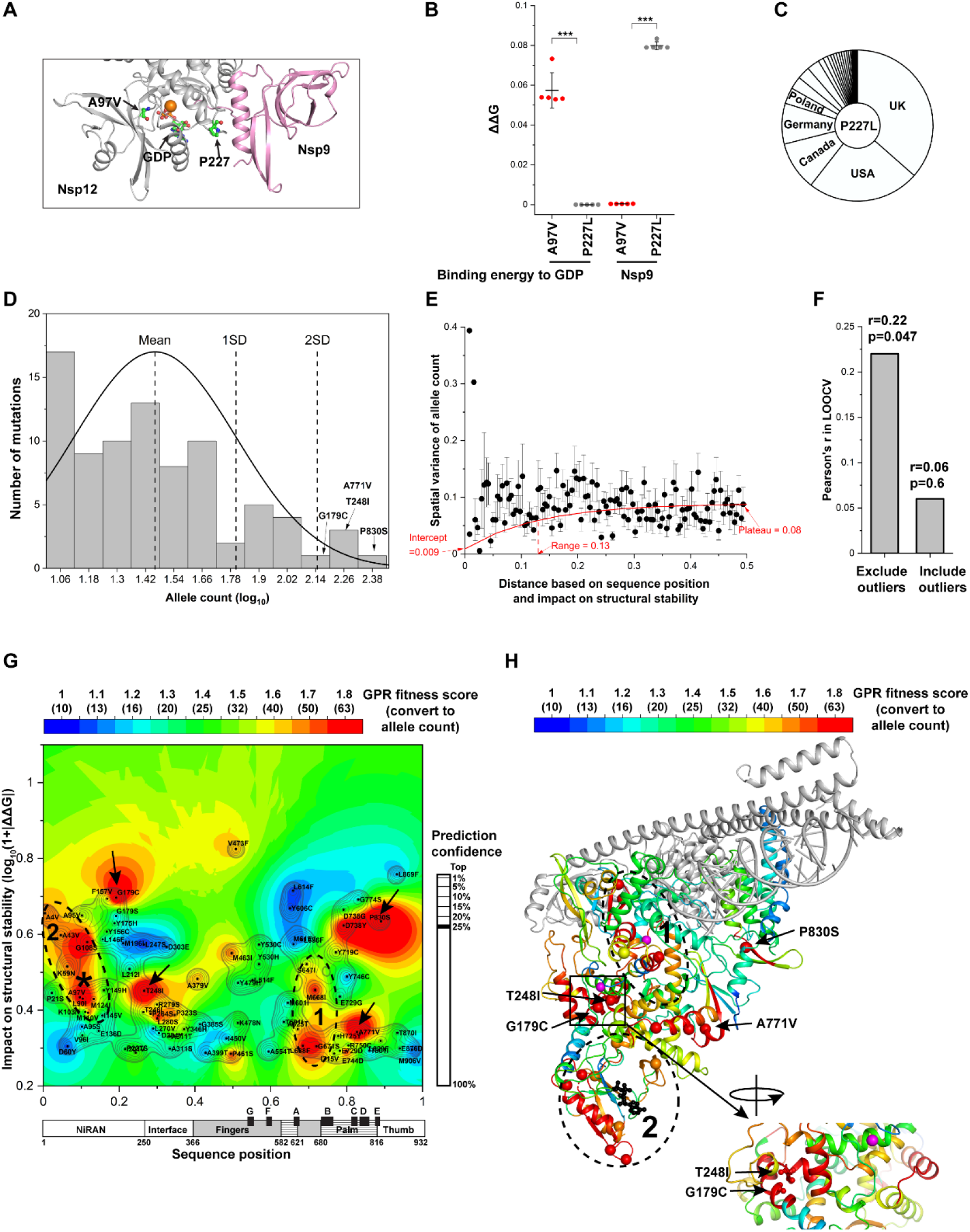
GPR model for nsp12 mutations found in the Alpha VOC. (**A**) P227 is located in the interface between NiRAN domain of nsp12 and nsp9. (**B**) P227L shows significant destabilizing signal for the binding energy to nsp9 when compared with A97V (Student’s t-test). (**C**) Country distribution of P227L. (**D**) Mutations that have significant stabilizing (ΔΔG>0.92 kcal/mol) or destabilizing (ΔΔG<-0.92 kcal/mol) impact with above ten allele counts are used to define the structural evolution of nsp12 in the Alpha VOC. G179C, T248I, A771V and P830S have allele counts that are above two standard deviations and are considered as ‘outlier mutations’ below. (**E**) Variogram analysis of the mutations excluding the outlier mutations G179C, T248I, A771V and P830S that are above two standard deviations. (**F**) Leave-one-out cross- validation of the GPR model with or without the outlier mutations. The GPR model excluding the outlier mutations achieves significant cross-validation results while the GPR models including the outliers do not, indicating that the outlier mutations are not clustered with other mutations with high GPR fitness scores. We used the GPR model excluding the outlier mutations in the main Fig. 3C to better illustrate the clustering effect of the mutations that drive the covariant fitness clusters. (**G**) GPR fitness landscape including the outlier mutations that are above two standard deviations (G179C, T248I, A771V and P830S). Consistent with the GPR model excluding the outlier mutations (Fig. 3C), we also observe covariant fitness cluster 1 and 2 in the fitness landscape including the outlier mutations. In addition, we observe that the four outlier mutations give high fitness scores to the regions surrounding them (**G**, black arrows). (**H**) Structural mapping the fitness landscape including the outlier mutations that are above two standard deviations. P830S is located isolated near RNA binding site. A771V is located at the connection region between cluster 1 and cluster 2. G179C and T248I are located on an interacting residue pair G179-T248, suggesting a critical residue-residue interaction in nsp12 for the fitness of the Alpha VOC.

**Figure S6.**
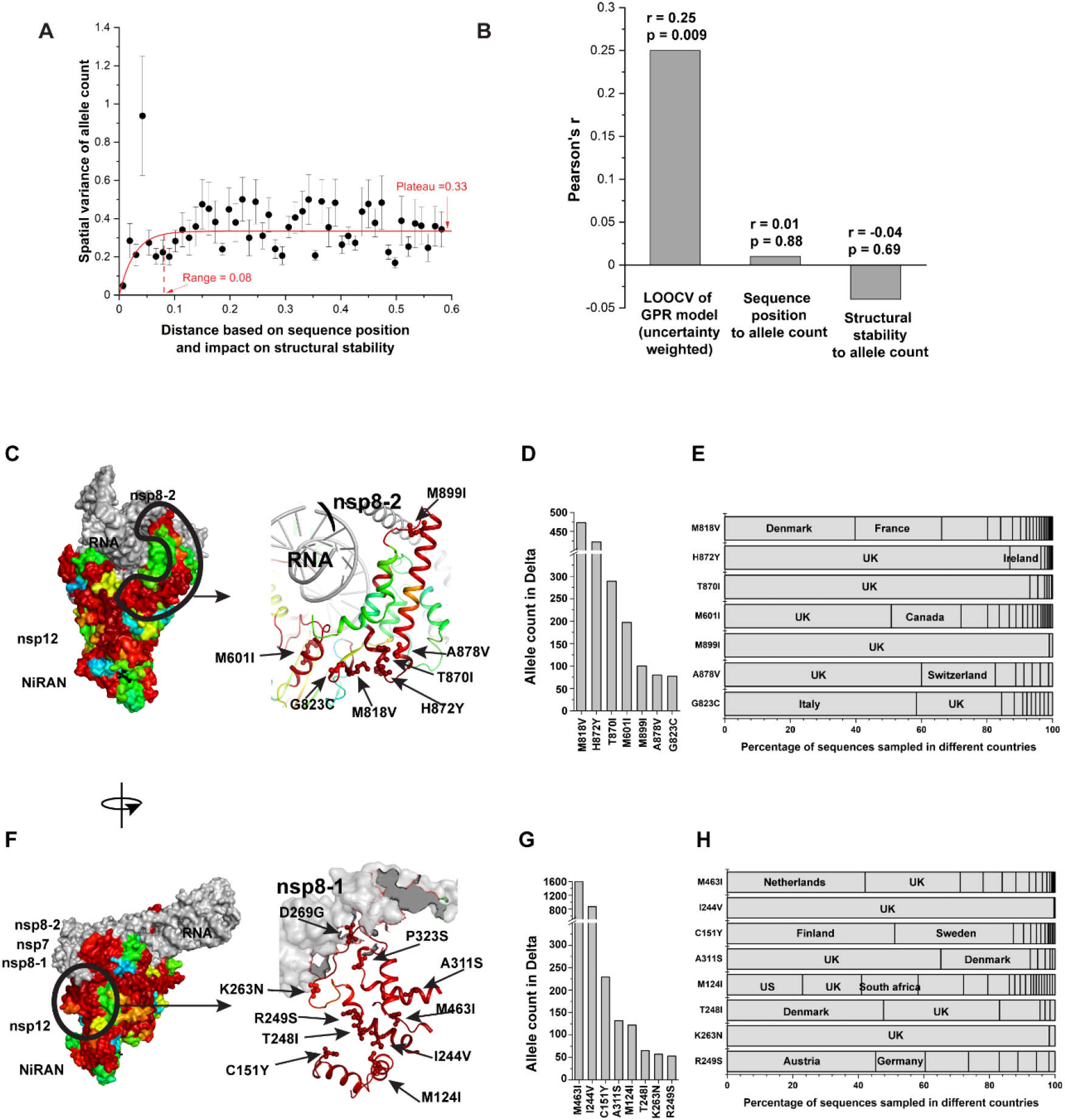
GPR model for nsp12 mutations found in the Delta VOC. (**A**) Variogram analysis of nsp12 mutations in Delta VOC that that have significant stabilizing (ΔΔG>0.92 kcal/mol) or destabilizing (ΔΔG<-0.92 kcal/mol) impact with above ten allele counts. (**C**) Leave-one-out cross- validation for the GPR model shown in Fig. 4E. Using the uncertainty generated from GPR model as weight (the weight is calculated as: ω=1/σ^2^, where σ^2^ is the GPR variance, see **Method**), the cross-validation yields Pearson’s r = 0.25 between actual allele count and predicted allele count with p = 0.009 (null hypothesis of r = 0; ANOVA test). There is no significant correlation between sequence position and allele count (Pearson’s r = 0.01, p = 0.88) or between mutation impact on protein stability and allele count (Pearson’s r = -0.04, p = 0.69), suggesting that there is additional relationship captured by GPR to generate the significant prediction for allele count (**B**). (**C-E**) Extension of the covariant fitness cluster of C487-H642-C645-C646 Zn^2+^ binding motif to the binding site between nsp12 and RNA/nsp8-2. The allele count and country distribution for each mutation are presented. The most frequent mutations in this region, M818V and H872Y are an interacting residue-residue pair (8 Å between C_α_ atoms). Whereas M818V evolved in Denmark and France, H872Y was largely circulating in the population of UK and Ireland. These results highlight that a similar structural feature adjustment can evolved independently in different countries to achieve improved fitness. (**F-H**) Residues with high GPR fitness scores linking NiRAN domain to the binding interface between nsp12 and nsp8-1. The allele count and country distribution for each mutation are presented. The two most frequent mutations in this region, M463I and I244V are within 8 Å separation of the C-alpha atoms. Interestingly, they show differential impact in different countries (Netherlands for M463I; UK for I244V), likely reflecting population specific features impacting the dynamics of spread.

